# A unique mechanism explaining the outstanding performance of a newly discovered doxycycline riboswitch

**DOI:** 10.1101/2025.11.11.687352

**Authors:** J. Hoetzel, A. Walbrun, M. Schäfer, T. Wang, A.G. Jørgensen, K. Stamatakis, V. Gunawan, O. Becker, L. Reichardt, L. Boettger, R.W. Bruckhoff, J. Kjems, J. Wachtveitl, M. Rief, B. Suess

## Abstract

Synthetic riboswitches offer a versatile and protein-independent solution for conditional gene regulation. They consist of a regulatory domain and an aptamer domain that binds a specific ligand, with their performance largely depending on the ability of the aptamer to control the regulatory domain. Expanding the range of synthetic riboswitches therefore requires the discovery and characterization of new regulatory aptamers. In the present study, we identified a doxycycline-binding aptamer with outstanding regulatory properties in both yeast and human cells which are based on a unique structural dynamic upon ligand binding. Single-molecule force spectroscopy revealed that doxycycline binding strongly stabilizes an intermediate aptamer conformation offering mechanistic insights into its function. The identification of the aptamer through a combination of parallel SELEX and subsequent *in vivo* screening in yeast, also provided valuable insights into selection dynamics and the first proof for the effectiveness of RNA Capture-SELEX in aptamer selection. Together, the presented data deepen our understanding of regulatory aptamer selection and functionality while adding a high-performing doxycycline-responsive aptamer to the synthetic biology toolbox.

## Introduction

Efficient and precise gene regulation is a fundamental prerequisite for many aspects of biological and medical research. Conditional gene expression enabled for example to study brain development, oncogenesis or viral infections in detail.^1–3^ Common tools used for this task are the TetON-system, utilizing different variants of the bacterial Tet-repressor, or the CRISPR-Cas System, using proteins of the Cas-family, to control gene expression on DNA or RNA level.^4–7^ Despite their excellent performance, they are suboptimal for a variety of applications that have to be either protein-independent or require the precise regulation on single transcript level.

Riboswitches offer a compact, versatile and protein-independent solution for the task of gene regulation and even beyond.^8^ Naturally found mainly in the 5’UTR of many bacteria, they exert control over transcription or translation of the mRNA they are located on.^9–11^ In most cases these riboswitches are responsive to metabolites functionally connected to the protein encoded by the controlled mRNA.^12–14^ Beside their prominent role in bacteria, a few examples have also been discovered in the mRNA of fungi, archaea and plants, leading to over 55 classes of natural riboswitches discovered to date.^15–17^ Meanwhile, this elegant concept of gene regulation has been adopted for the engineering of synthetic riboswitches. They are constructed of an aptamer domain, sensing a specific ligand, and a regulatory domain which allows for gene regulation.^18^ Due to their simple yet effective architecture, synthetic riboswitches have been successfully applied outside of their bacterial origin for regulatory purposes to all domains of life. Like their natural counterpart, synthetic riboswitches operate through structural changes that are induced by the binding of their ligand to the aptamer domain.^19–22^ These changes in turn are transmitted to the regulatory domain and ultimately lead to regulation of gene expression. Due to their modular nature, synthetic riboswitches have been engineered to operate in a vast number of different functional mechanisms. Depending on the regulatory domain used, this allows for different modes of gene regulation in a wide range of different species including for example splice-control in human cells,^23^ blocking of ribosomal scanning in yeast,^24^ control of RBS accessibility in bacteria and archaea,^25,26^ control of ribozyme cleavage in *C. elegans*^27^ or control of polyadenylation in mice.^28^ Despite their differences, all aforementioned synthetic riboswitches share the characteristic of utilizing an aptamer as their sensor domain that was selected *in vitro* through Systematic Evolution of Ligands by Exponential Enrichment (SELEX).^29,30^ Developed for the purpose of selecting nucleic acid binding motifs *de novo*, SELEX opens the possibility to select aptamers binding to virtually any given ligand. This creates the possibility to engineer riboswitches using the same regulatory domain but aptamers responsive to different ligands. In turn, this enables orthogonal gene regulation that allows to independently control multiple riboswitches in the same cell. Their versatility has also led to applications beyond mere gene expression regulation like the intracellular detection of metabolites by fluorogenic readout or *in vitro* biosensing.^31–33^

Regardless of the final application of a riboswitch, a decisive element of their functionality is the utilized aptamer domain. Although several hundred RNA aptamers have been selected since the invention of SELEX only a handful have been found suited for the engineering of riboswitches. Because of the structural changes necessary to exert regulation over the regulatory domain of the riboswitch, only aptamers that are able to undergo these changes upon ligand binding can be used for riboswitch engineering.^34^ These structural changes are accompanied by a high affinity to the ligand and a distinct 2-step binding mechanism, observed in regulatory aptamers like the tetracycline, neomycin or ciprofloxacin binding aptamers.^35–37^ To facilitate the selection of novel aptamers which meet these criteria, two main factors in the selection have been adjusted: the SELEX method itself and subsequent screenings to identify suited candidates. With the selection step being a decisive factor for the final applicability of the selected aptamers, SELEX methods like Capture-SELEX, *in vivo* SELEX or ultraSELEX have been developed to increase the selection efficiency of aptamers fit for their intended application.^38–40^ Among the specialized SELEX methods, RNA Capture-SELEX was developed for the effective selection of RNA aptamers undergoing conformational changes upon ligand binding.^41^ In contrast to conventional SELEX methods, the RNA pool is coupled to beads through hybridization to a DNA oligonucleotide while the ligand is free in solution (conventional SELEX methods require immobilization of the ligand) (Fig. 1b). In this setup, only candidates able to undergo structural changes upon ligand binding and detach from the DNA oligonucleotide are subsequently selected. Furthermore, previous attempts of selections have also shown that a subsequent screening is crucial for the effective identification of suited candidates.^42^ When aiming to engineer riboswitches, an *in vivo* screening in *Saccharomyces cerevisiae* has proven to be an efficient way to identify aptamer candidates with promising characteristics.^42,43^

**Figure 1:**
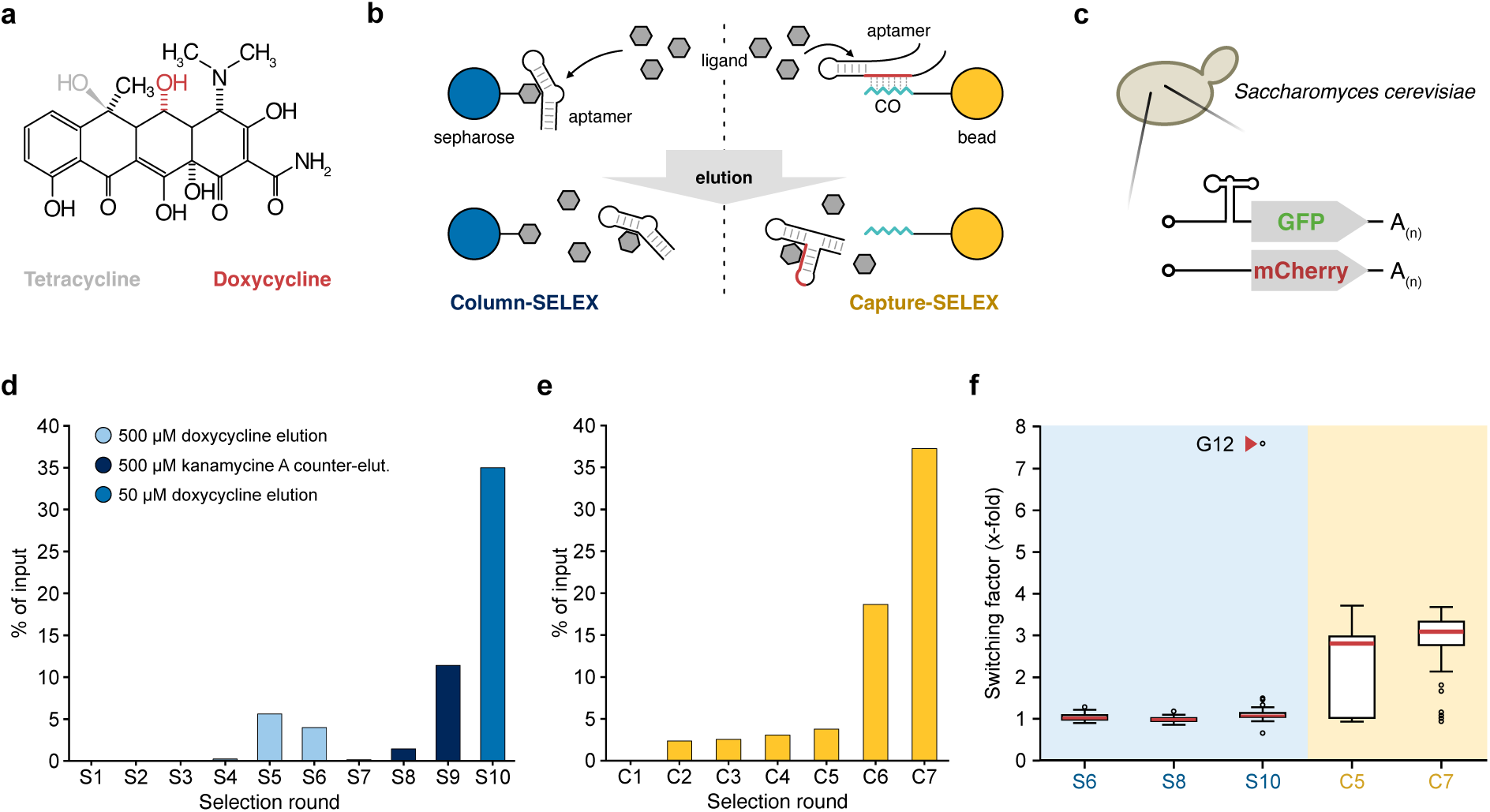
*In vitro* selection and subsequent *in vivo* screening of doxycycline-binding aptamers **a)** Natta projection of the shared structure of tetracycline and doxycycline. The two tetracyclines differ in the position of a single hydroxyl group. Positions of the distinctive hydroxyl group in doxycycline (red) and tetracycline (grey) is indicated. **b**) Schematic depiction of the selection step in conventional and Capture-SELEX. The target ligand (grey) is immobilized on a column material (blue) in the conventional SELEX. During Capture-SELEX, the RNA pool is immobilized by hybridization of a docking sequence (red) to the capture-oligonucleotide (turquoise) which is coupled to a paramagnetic bead (yellow). The elution of bound RNA sequences is performed in the conventional SELEX by washing with high concentration of the target ligand, whereas in the Capture-SELEX the RNA that undergo conformational changes upon ligand binding detach from the capture-oligonucleotide and are subsequently eluted. **c**) Schema of the translation initiation control mechanism in *S. cerevisiae* used for *in vivo* screening and aptamer testing. A double reporter plasmid brought into yeast, leading to transcription of a GFP mRNA containing the aptamer candidate in the 5’UTR and a mCherry mRNA with no aptamer structure. **d**) Progress of the conventional SELEX depicted in eluted RNA fraction in percent relative to the input. The different selection parameters over the course of selection are indicated by colour: elution with 500 µM doxycycline (light blue), additional counter-selection with 500 µM kanamycin A (dark blue) and elution with reduce doxycycline concentration of 50 µM (navy blue). **e**) Progress of the Capture-SELEX depicted in eluted RNA fraction in percent relative to the input. **f**) Results of the *in vivo* screening in *S. cerevisiae* of the pools obtained from conventional SELEX (blue background) and Capture-SELEX (yellow background). Plotted on the x-Axis are the tested selection rounds and their switching behaviour is indicated on the y-Axis in x-fold dynamic range. The red line in the box plots depict the median switching value of the pool and circles depict outlier from the population behaviour. The aptamer G12 is highlighted by a red arrowhead in the pool S10.

In the present study, we performed *de novo* selection for a new regulatory aptamer binding doxycycline applying two different *in vitro* selection methods to compare selection outcome of Capture-SELEX and a conventional SELEX approach. Doxycycline was chosen as the ligand for its low toxicity and its already widespread presence for the actuation of Tet-based systems.^7^ After successful selection, we gained valuable insights into the selection process by NGS analysis, providing the first evidence for the high efficiency of Capture-SELEX for the selection of regulatory RNA aptamers. The discovered regulatory aptamer G12 was identified by *in vivo* screening in yeast and possesses -to the best of our knowledge-the best regulatory capability of a regulatory aptamer described to date. Comprehensive characterization by mutational studies, structural analysis and binding dynamic studies revealed a unique structural dynamic upon ligand binding with the strong stabilisation of an intermediate conformation of the aptamer, driving the regulatory potential of G12. Overall, we gained exciting insights into SELEX and molecular dynamics within regulatory aptamers while providing a thoroughly characterized, high performing aptamer ready to be utilized for synthetic riboswitch engineering.

## Results

### Discovery of doxycycline-binding aptamers with regulatory potential by two *in vitro* selections and subsequent *in vivo* screening

Based on our hypothesis that Capture-SELEX is the most effective method for the selection of aptamers undergoing conformational changes upon ligand binding, we started both *in vitro* selections with the identical starting library to compare the selection outcome from both selections (Fig. S1a).

Beginning with the conventional column-based SELEX, we obtained the first signs of enrichment of binding to doxycycline after 5 rounds (Fig. 1d, Table S1). Increase in washing steps and introduction of a counter-selection with kanamycin A to remove weak or unspecific binding sequences, led to a decrease in eluted RNA. The elution fraction recovered until round S9 and even further increased in round S10 when counter-selection was left out for an elution step with reduced doxycycline concentration. With 35 % of the RNA input eluted in round S10, the pool showed strong signs of an enrichment of target binding sequences. Capture-SELEX was performed as previously described starting with the identical library as used for the conventional SELEX and 100 µM of doxycycline for the elution of candidates.^41^ After six rounds of selection an increased elution of RNA was observed and further increased in round C7 (Fig. 1e & Table S2).

Pools of round S6, S8 and S10 from the conventional SELEX and round C5 and C7 from the Capture-SELEX were used for *in vivo* screening in *Saccharomyces cerevisiae*.^44^ The aptamer candidates were cloned into the 5’UTR of a GFP gene, where they can act as a roadblock to the pre-initiation complex of the ribosome and thereby regulate translation of GFP mRNA. From the same plasmid mCherry is constitutively expressed to serve for normalisation of GFP expression. First, we pre-selected the sequences for sufficient expression levels in the absence of doxycycline. With progressing selection rounds from the conventional SELEX, we observed a reduction in the preselected fraction (Fig. S1c). This was not observed for the rounds obtained from Capture-SELEX (Fig. S1d) and implies a difference in enriched sequences over the selection progress of both SELEX methods. Among the pre-selected candidates from the conventional SELEX no noticeable switching behaviour was observed upon addition of doxycycline, except for a single candidate with a 7x dynamic range (Fig. 1f). The pools from the Capture-SELEX showed an overall median dynamic range of around 3x in round C5 and C7 (Fig. 1f), indicating a great fraction of sequences with prominent switching behaviour in these pools. This presented a striking difference in selection outcome between the two selections in relation to sequences undergoing conformational change upon ligand binding. Sequencing of the candidates with regulatory capability revealed one aptamer sequence from the conventional SELEX named G12 and one aptamer causing the shifting in the pool from the Capture-SELEX named C01.

### Next-generation sequencing reveals differences in selection dynamics between SELEX methods

The stark difference in number of regulatory aptamers found in the rounds from the two SELEX methods, raised the question how the dynamics within the selections shaped the composition of sequences. The selection mechanism in a SELEX method heavily influences the characteristics of the selected sequence, like in this case their regulatory potential.^34^ To shed light on the selection dynamics, all selection rounds were sequenced by next-generation sequencing for bioinformatic analysis. From the obtained sequences the ten most abundant motifs in each round of 6 to 12 nucleotides length were identified by MEME, mapped by FIMO for similarity and collapsed to motif clades (Fig S2a-S2d). The resulting motif clades were then tracked in their abundance over the course of their respective SELEX. A total of 49 motif clades were identified for the pools from the conventional SELEX. The composition of motif clade abundance changed over the course of selection rounds and with changing selection parameters (Fig. 2a). Especially after the introduction of counter-selection, the composition of the pool changed with some clades being reduced in abundance while others increased. In contrast, the reduction in doxycycline concentration for elution in round 10 caused no comparable effect, leaving the motif clade composition between round 9 and 10 mostly unchanged. The composition of motif clades behaved different for the Capture-SELEX, with 22 motif clades identified and a fast enrichment of few clades within 7 rounds of selection (Fig. 2b). Within the first five rounds of selection, the diversity of motif clades present in the pool was quickly reduced to seven motif clades with considerable abundance. Between selection round 5 and 7, six motif clades (clade 2, 3, 4, 10, 16 & 20) accumulated in a dominant way. When checking the origin of these clades, it turned out that three of these motif clades can be attributed to parts of the aptamer C01 (Fig. S2e). In comparison, none of the dominant motif clades from conventional SELEX could be attributed to G12 or C01. To understand the enrichment of the two aptamers, we followed their abundance in the pools over both selections. During conventional SELEX, C01 fluctuated in abundance throughout the selection starting among the Top25 sequences in the first rounds and then falling out of the Top100 sequences while still increasing in read count (Fig. 2c). G12 started very low and did not increase in abundance until round 8. Both aptamers decreased in abundance in round 10 with the reduced doxycycline concentration in the elution step. In the Capture-SELEX, C01 was found after round C1 among the Top10 sequences and became the dominant sequence from round 4 on and continuously enriched until the final round 7 (Fig. 2d). In contrast, G12 was very low in abundance, never among the Top100 sequences and showed only signs of enrichment in the last three rounds of selection. To understand this behaviour of G12, we tested its capability to couple to the capture-oligonucleotide in the Capture-SELEX setup and found out that it was unable to effectively hybridize to the capture-oligonucleotide (Fig. S1e).

**Figure 2:**
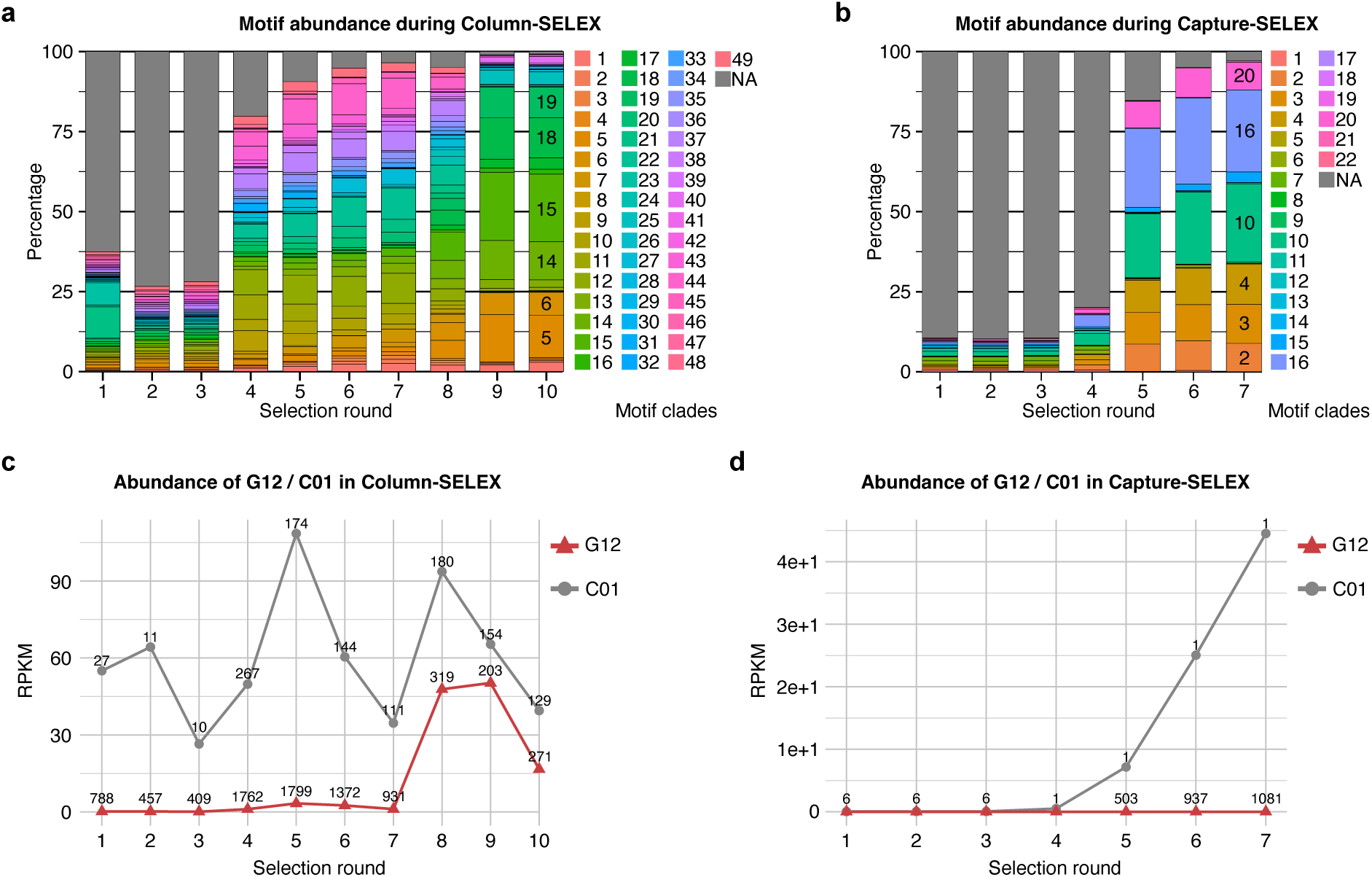
Investigating selection dynamics via NGS analysis. **a**) Tracking of the most abundant motif clades over the course of the conventional SELEX selection. The different motif clades are indicated by colour and the plotted by their relative abundance in percent of all analysed reads. Dominant clades in the last selection rounds are indicated by their clade number. **b**) Tracking of the most abundant motif clades over the course of the Capture-SELEX selection. **c**) Tracking of the two aptamer candidates G12 (red) and C01 (grey) over the course of the conventional SELEX. For each round the abundance of the candidates is indicated normalized in reads per million (RPKM) on the y-axis and the relative abundance as position among all sequences (indicated over data point). **d**) Tracking of the two aptamer candidates G12 (red) and C01 (grey) over the course of the Capture-SELEX.

### Characterization and optimization of the new doxycycline-binding aptamer G12

We decided to continue characterisation of G12 due to its outstanding regulatory behaviour. The folding prediction of G12 proposed a T-shaped secondary structure with a closing stem P1, transitioning into the J1-3 three-way junction.^45^ Helices P2 and P3 ending in a twelve and six nucleotide long loop, respectively (Fig. 3a). We performed mutational studies and chemical probing to proof this predicted structure. First rational mutations stabilising the P1 stem by exchanging nucleotides leading to perfect base pairing (M1, M2) were tested and maintained regulatory activity while showing reduced expression levels (Fig. S3a, S3d). Truncation of the P1 stem to seven base pairs (M3) increased the dynamic range without lowering expression levels in the absence of doxycycline. Replacing the sequence of the L3 loop (M4, M5) led to complete loss of regulatory activity. To further validate the importance of the P3 stem, we disrupted the stem by two consecutive mutations (M9) leading to loss of regulation. The introduction of two additional mutations (M10) restoring base pairing recovered the activity. We continued characterisation based on M3 since it was the best performing candidate (Fig. S3d). Flipping the sequence of the P3 stem (M11) led to performance comparable to M3 (Fig S3b, S3e). The insertion of one additional base pair (M12) reduced dynamic range and the insertion of two base pairs (M13) led to complete loss of regulatory behaviour. These observations indicate a structural function of the P3 stem, independent of its sequence (Fig S3b, S3e).

**Figure 3:**
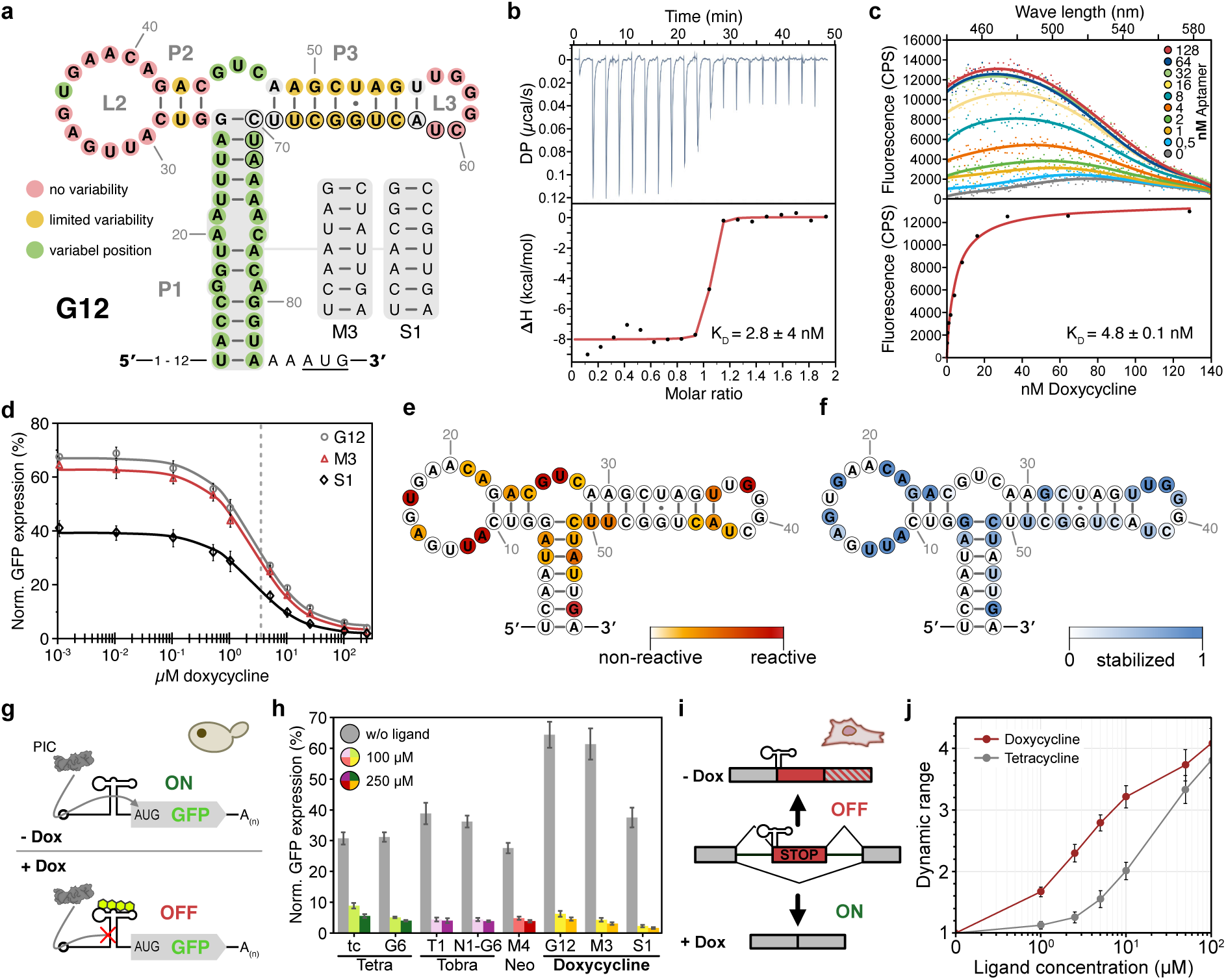
Characterization of the doxycycline-binding aptamer G12. **a**) Depiction of the folding prediction of G12 as found in the *in vivo* screening. The colour coding represents the summary of the findings from the rational and randomized mutational studies, with red indicating positions with no mutation tolerance, yellow indicating limited mutation tolerance and green indicating positions that allows free nucleotide variability. Alternative P1 stems as used in the variants M10 and S1 are shown beside the aptamer. Nucleotide positions in the P3 with a black circle represent the docking sequence as found in the Capture-SELEX pool. **b**) Representative ITC results of G12 showing the titration curve and thermogram. **c**) Representative FTS results of G12 showing the fluorescence curves of different aptamer concentrations and the curve fit over the experimental data. **d**) Dose response curve of G12, M10 and S1 in yeast regulating GFP expression through the roadblock mechanism. Expression levels are shown over varying doxycycline concentrations, indicated on the x-axis, in percent normalized and relative to a consecutive GFP expressing positive control (pCBB05), indicated on the y-axis. Experimental data is represented by the data points, and the fitted curve is shown in corresponding colour. The EC50 of all three constructs was close to 3 µM as indicated by the grey, dashed line. Error bars depict the standard deviation of two independent measurements with 3 biological replicates. **e**) Chemical probing of M3 by SHAPE-MaP in the absence of doxycycline. Reactivity of each nucleotide position is indicated by colour (red = highest reactivity) in the 2D representation of the aptamer. **f**) Stabilization of nucleotide positions as calculated from SHAPE-MaP results of M3 with and without doxycycline and mapped in colour to the 2D representation of the aptamer. Intensity of colour (blue) indicates reduced reactivity of a nucleotide position, implying increased position, stability upon binding of doxycycline. **g**) Schema of translation initiation control in yeast through the roadblock mechanism. Translation of the reporter gene (GFP) is controlled by the stability of the aptamer placed in the 5’UTR of the gene. In absence of its ligand, the aptamer is in a rather loose state, allowing the ribosomal pre-initiation complex (PIC) to pass. Upon binding its ligand, the aptamer stabilises and blocks the PIC from passing, ultimately supressing translation by keeping the PIC from reaching the start codon. **h**) Comparative measurement of different regulatory aptamers utilizing regulating translation through the roadblock mechanism. Expression levels are shown in percent normalized and relative to the positive control. Grey bars are showing the expression level of the constructs in the absence of their ligand; light coloured bars show the expression level in the presence of 100 µM ligand concentration and darker coloured bars show the expression levels at 250 µM ligand concentration. Error bars depict the standard deviation of two independent measurements with 3 biological replicates. **i**) Schema of the mechanism controlling mRNA processing through 3’ splice site accessibility. The synthetic exon (red) containing premature stop codons is spliced into the mRNA in absence of doxycycline, due to the accessibility of the splice site to the spliceosome, leading to premature termination of translation. In presence of doxycycline, the splice site is masked and the synthetic exon sliced out from the mRNA. **j**) Dynamic range of reporter gene expression at different concentrations of ligands of the doxycycline-binding riboswitch (red) and the tetracycline-binding riboswitch (grey) in HeLa cells. The riboswitch construct containing G12 shows a faster response to lower concentrations of its ligand and an overall higher dynamic range when compared with the tetracycline riboswitch. Data points and standard errors were calculated from three independent measurements in triplicates.

Exchange of the three nucleotides in the J1-3 three-way junction (M14) led to comparable behaviour to M3, implying no importance of this sequence for the function of the aptamer (Fig. S3b, S3e). All mutations altering the sequence of the P2 stem (M15, M16, M18, M19) led to great reduction or loss of regulatory performance of the aptamer, except for the flipping of the middle A-U base pair (M17) which maintained similar overall behaviour to M3 (Fig. S3b, S3e, S3f). Any exchange of a triplet in the L2 loop into CAA (M20-M23) caused complete loss of regulatory activity (Fig. S3b, S3f). These results indicate importance of the sequence of the two loops L2 and L3 and mainly structural function of the P2, J1-3 and P3 region.

For a detailed analysis on single nucleotide level, we performed a random mutation screening. Random single mutations were introduced from G7 to C51 in the M3 aptamer, screened for their regulatory activity and analysed by NGS. The results showed mostly loss of function upon insertion of any single nucleotide mutation (Fig. S3c). Only five positions were found tolerating mutations, being a single position in the L2 loop (U17), the three nucleotides in the J1-3 junction (G26, U27, C28) and the exchange of the position G46 in the P3 stem for an adenosine. These findings confirm the results of the rational mutations, indicating importance of the sequences in the L2 and L3 loops. Moreover, it supports the scaffolding function of the stems P1, P2 and P3. The results are summarized in Figure 3a.

Based on our experience with synthetic riboswitch engineering the composition of the closing stem P1 highly influences the performance of a riboswitch.^24,46^ We created several stem variants based on M3 with varying stem stabilities (S1-S5, Fig S3b). Testing in yeast resulted in S1 as the best performing candidate (Fig. S3g).

Next, we determined the KD of the aptamer variants G12, M3 and S1 by isothermal titration calorimetry (ITC) and fluorescence titration spectroscopy (FTS, Fig. 3b & 3c, Fig. S4a-i). Equal affinities to doxycycline of around 3 nM measured for all three aptamers indicate that the improvement of regulatory behaviour was not caused by improved ligand binding. Importantly, none of the three variants showed signs of binding to tetracycline, a close derivative only differing in one hydroxyl group from doxycycline (Fig 1a, Fig. S5a-S5f). Further, we determined a comparable EC50 value of 2,7 µM for all three variants responding to doxycycline in yeast, confirming identical response to the ligand and underlining effects of the scaffolding role of the P1 stem for regulation (Fig 3 d, Table S4).

Finally, we performed SHAPE-MaP to confirm our previous structural insights.^47,48^ Nucleotide reactivity of the variant M3 was examined in the absence and presence of doxycycline. Surprisingly, we observed unexpected low overall reactivity, suggesting a pre-structured RNA folding (Fig. S6a & S6b). In line with our mutational studies, we observed a high flexibility in the three-way junction J1-3. Additional flexibility was observed in the loops L2 and L3. However, only few nucleotides in these regions showed considerable reactivity, indicating involvement of the loops in tertiary structure formation. The high reactivity of positions U17, G26 and U27 correspond well with the findings of the mutational studies (Fig. S3c). Strikingly, vast parts of the aptamer are stabilised upon binding to doxycycline (Fig. 3e, Fig. S6e, S6f). Again, in line with the mutation studies, most of the stabilised positions are located in the L2 and L3 loop, underlining importance of these structures. Furthermore, the upper part of the closing P1 stem is stabilised by binding to doxycycline.

### Application *in vivo* reveals great regulatory potential of G12 in yeast and human cells

The new doxycycline aptamer clearly outperforms previously described synthetic riboswitches. In a comparative measurement of controlling translation initiation in yeast, G12 and its variants M3 and S1 were compared with the well-characterized and commonly used tetracycline aptamer,^49^ the neomycin aptamer^50^ and the new and high-performing tobramycin aptamer.^43^ For the tested ligand concentrations of 100 and 250 µM, all three versions of the new doxycycline-binding aptamer showed the highest dynamic range (Fig. 3h & Table S5). The observed high dynamic range of the G12 variants can be largely attributed to a relatively high expression level in the absence of its ligand with a comparable or even lower expression levels in the presence of the ligand, when compared with the other aptamers. Particularly, the variant S1 possessed the best regulatory behaviour overall with an expression level of 38%, and a reduction down to 1,5% at 250 µM doxycycline, resulting in a dynamic range of 25-fold. Since the regulation through the roadblock mechanism in yeast is greatly defined by the structural stability of the aptamer, the switching behaviour of the doxycycline-binding aptamer suggests a substantial structural dynamic between the ligand-bound and unbound state.

For gene expression control in human cells, we applied the new doxycycline-binding aptamer for regulation of mRNA processing by controlling 3’ splice site accessibility.^23^ The G12 aptamer was placed on the original P1 stem incorporating the 3’ splice site to control the recognition of this site by the spliceosome. In the absence of doxycycline, the P1 allows recognition of the 3’ splice site by the spliceosome, leading to inclusion of a synthetic exon containing multiple premature stop codons and ultimately perturbing translation of the target gene (Fig. 3g). Through conformational changes of the aptamer upon binding of doxycycline, the P1 stabilizes and sequesters the 3’ splice site, leading to exclusion of the synthetic exon. Although the riboswitch context was originally optimized for the tetracycline aptamer, the new doxycycline aptamer showed outstanding regulatory performance without any further adaptations. In comparison to the tetracycline aptamer, G12 possessed regulatory response lower concentrations of its ligand than the tetracycline aptamer and an overall higher dynamic range (Fig. 3h & 3i).

### G12 possess a unique intermediate conformation stabilised by binding to doxycycline and independent of the P1 stem

With the outstanding regulatory performance of G12 compared to other regulatory aptamers, we expected a pronounced dependence of aptamer stability on ligand binding. To quantify these changes, we performed single-molecule force spectroscopy, which provided mechanistic insights into the structural stability of G12. The aptamer was immobilized using DNA linkers between two beads held by laser foci (Fig. 4a; for details see Materials and Methods and ^51^). Our setup allows the application and quantification of pulling forces on the RNA structure while simultaneously enabling the transfer of the aptamer between microfluidic channels containing different solution conditions (Fig. 4b).

**Figure 4:**
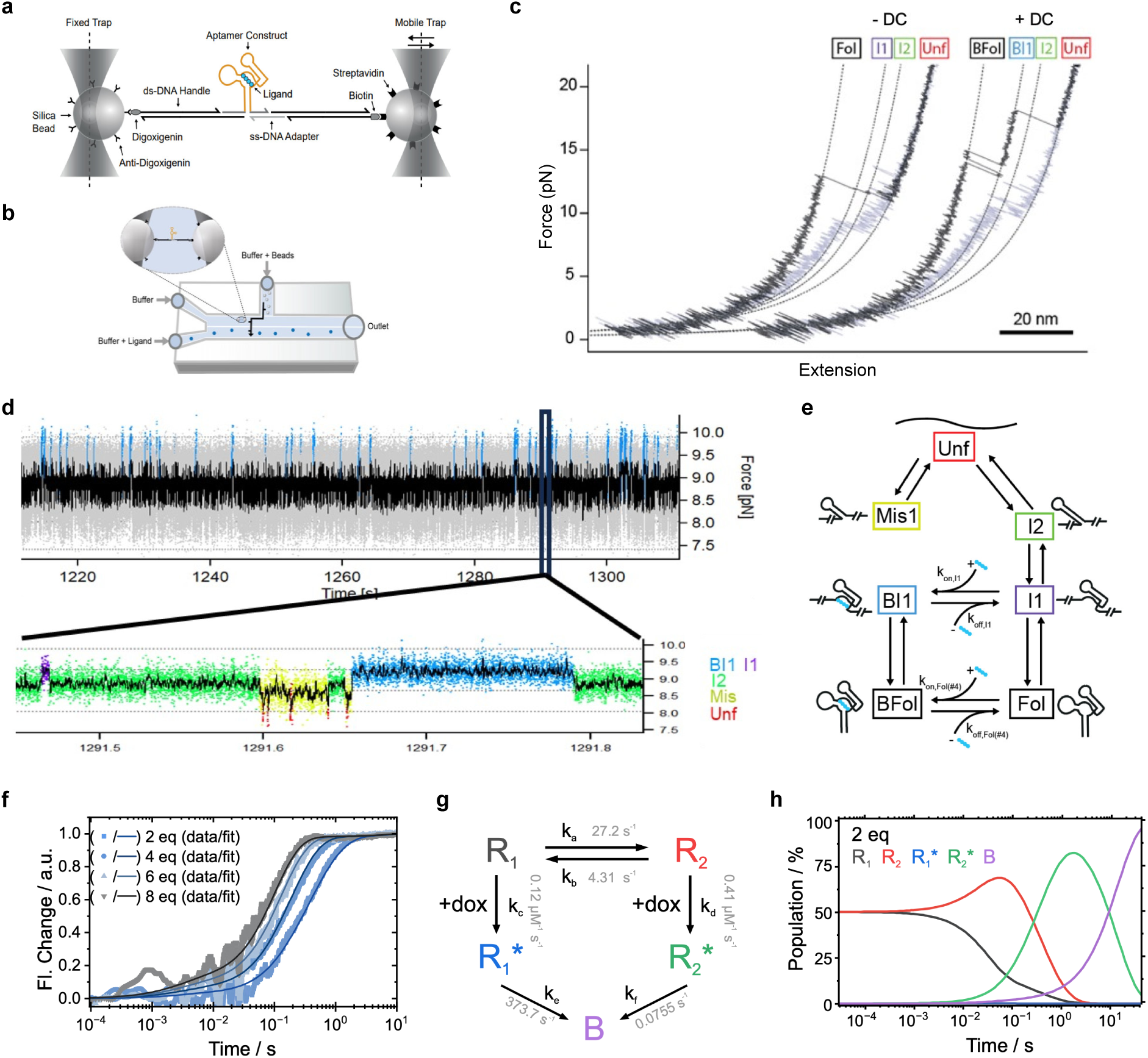
Investigation of structural and binding dynamics of G12. **a**) Schematic depiction of the optical tweezer setup with the aptamer immobilized by hybridization to DNA handles between two silica beads. **b**) Schema of the microfluidic setup used for the molecular tweezers using different channels to change measurement environment and parameters. **c**) Representative curves of force-ramp measurements of the short P1 stem variant of G12 with and without doxycycline. Plotted on the x-axis is the extension of the setup and the resulting resistance in force (pN) is plotted on the y-axis. The identified intermediate structures are indicated above the curves. **d**) Representative trace of a constant distance measurement of the G12 variant without a P1 stem. Plotted on the x-axis is the time in seconds and on the y-axis the observed force in piconewton. The upper trace represents a full measurement with an extended timeframe, while the lower trace is a zoomed-in section as indicated. The identified intermediate states bound intermediate 1 (blue), intermediate 1 (purple), intermediate 2 (green), misfolded (yellow) and unfolded (red) are indicated by colour. **e**) Schematic of the folding and transition dynamics of G12 between intermediate conformations, ranging from completely unfolded to fully folded and bound to doxycycline. Transitions from an unbound to a bound state include the kON and kOFF values of the corresponding transition. **f**) Experimental data from stopped flow experiments and the fitted curves from the best fitting model describing the binding dynamic. The time observing the fluorescence is plotted on the x-axis with the observed change in fluorescence plotted on the y-axis. The experimentally measured data at different equivalents of aptamer to doxycycline are shown as data points and the fitted curve from the model as a coloured line. **g**) Schema representing the different conformations needed to explain the observed binding dynamic. The aptamer has two unbound conformations (R1, R2), two intermediate bound conformations (R1*, R2*) and one fully bound conformation (B). The unbound conformations are in equilibrium with a preference to fold into R2. Association rates are indicated on their respective steps. **h**) Modelling of the populations of the different over time and binding to doxycycline, best explaining the observed binding behaviour. The populations were plotted in the y-axis as percent of all sequences over time plotted on the x-axis. The shown graphs were calculated for 2 equivalents of aptamer to doxycycline and were started for 1:1 ration between the unbound conformations R1 and R2.

In a first set of experiments, we probed the structural stability of a G12 variant with a 5-base-pair-long P1 stem (short stem). In force-ramp experiments (Fig. 4c), we observed that, in the absence of doxycycline, the aptamer unfolds in a major peak at around 13 pN, followed by two short-lived on-pathway intermediates (I1 and I2). Representative selections of traces are shown in Fig. S8e. Upon addition of doxycycline, I1 is significantly stabilized and unfolds at even higher forces than the main peak, whose unfolding force remains unchanged (see force histograms in Fig. S7d & e). The strong stabilization of I1 enables repeated refolding to the folded state, resulting in multiple transitions between Fol and I1. After unfolding, the construct folds back into the native state (grey traces), though considerable hysteresis is observed. Force ramp measurements provide precise estimates of the unfolded portions of the intermediates through contour length fits (dashed lines), allowing us to propose coarse structural models for the various unfolding intermediates (see table S6 and Materials and Methods). We find that the length gains are consistent with a model in which the full P1 stem is unfolded in I1, and unfolding has proceeded until the P3 stem in I2. These data suggest that P1 is not critical for doxycycline binding, since the magnitude of the first peak does not depend on it.

We next examined the influence of P1 stem length by measuring two additional constructs: one with an extended P1 stem (long stem, 12 base pairs) and one lacking P1 entirely (no stem). Consistent with our expectation, the longer P1 stem exhibits a greater contour length gain and a slightly higher unfolding force, indicating overall increased stability (Fig. S8f). Similar to the short-stem construct, P1 unfolding forces are independent of doxycycline binding, and the stabilization of I1 by doxycycline is identical to that observed for the short-stem construct. Interestingly, the no-stem construct also displayed doxycycline-dependent stabilization of I1, identical to both of the other constructs, supporting the conclusion that the essential determinants for doxycycline binding remain intact even in the absence of P1.

As a reference, we measured the well-characterized tetracycline aptamer. Unlike G12, this aptamer unfolds in a single major cooperative peak, both in the presence and absence of tetracycline (Fig. SS7). This suggests that key determinants of tetracycline binding are also located within the P1 stem. Unfolding force histograms show higher forces in the presence of tetracycline, consistent with an energetic stabilization upon ligand binding. In direct comparison, these results highlight a distinct binding mechanism for G12, in which most of the interaction is mediated by an intermediate structure that does not involve P1.

To extract the dynamics and energetics of doxycycline binding to G12 in real time, we applied a constant force bias to a single molecule and observed near-equilibrium transitions between the various folding intermediates in the presence of doxycycline. Using the no-stem construct enabled us to monitor a single molecule for hundreds of seconds, capturing numerous binding events (Fig. 4D, BI1 blue states). A zoom into the trace reveals the known intermediates I1 (purple), BI1 (blue), I2 (green) and Unf (red), which were assigned by Hidden Markov Modeling (HMM) (see SI and ^52^ for more details). Despite occurring at the same force level due to structural similarity, BI1 could be distinguished from I1 by its markedly longer lifetime, reflecting stabilization upon doxycycline binding. We also identified a new state populated exclusively from the unfolded state (Unf), which we attribute to a misfolded conformation (Mis, grey). Such misfolded states are common in RNA folding and typically arise from incorrect base pairing from the fully unfolded state.^53^

A detailed kinetic analysis of the doxycycline-bound (blue) states provided on- and off-rates for binding to the no-stem construct (see Fig. S9b & c and Table S7; for details see Materials and Methods). We find that binding occurs exclusively to I1, as the measured on-rates correlate precisely with the probability of I1 population (Fig. S9). This strongly supports conformational capture as the mode of binding and implies that, within I1, the necessary tertiary contacts for doxycycline recognition are already formed. Interestingly, the observed off-rate of 6.0 s^-1^ appears relatively fast, and the calculated affinity of 3.8 µM is substantially weaker than the affinity of G12 measured by isothermal titration calorimetry and fluorescence titration spectroscopy.

We hence tested the influence of the P1 stem on binding kinetics with the short-stem construct. Unlike with the no-stem construct, similar equilibrium measurements were not possible, because, at the relevant forces, the short-stem construct will almost exclusively populate BFol. To overcome this, we developed a jump protocol where we moved the molecule between channels with and without doxycycline, allowing binding and unbinding rates to be measured at near-zero force (see Fig. S9 and Materials and Methods). These experiments revealed that the presence of P1 drastically decreases the off-rate (0.0055 s^-1^) while only marginally reducing the on-rate (0.42 s^-1^ µM^-1^) (Fig. S9g, Table S7). The resulting affinity of 13.1 nM closely matches the G12 affinites determined by isothermal titration calorimetry and fluorescence titration spectroscopy.

### G12 binds doxycycline in a two-step mechanism in two paths

For gaining insights into the aptamers binding dynamics, stopped-flow spectroscopy (SF) was performed utilizing the intrinsic fluorescence of doxycycline caused by binding to the aptamer. Four measurements with increasing ligand concentrations (2, 4, 6 & 8 equivalents) were conducted (Fig. 4f). The fluorescence increased with ligand binding and the binding process accelerated from 3 s to 1 s with higher equivalents of ligand. The binding mechanism of doxycycline to G12 was analysed with a kinetic modelling (KM) approach. Starting with a simple one-step binding process, the model was progressively refined by incorporating additional conformational states, intermediate steps and reverse reactions (Table S8). The comprehensive KM analysis is described in detail in the Supplementary Information. The best model of the KM depicted revealed two initial conformations of ligand free aptamer in equilibrium (Fig. 4g, Table S9). Both conformations (R1 and R2) bind doxycycline via a two-step binding mechanism, involving an initial ligand-aptamer association followed by a ligand independent conformational rearrangement. The population plot based on this model indicates that binding primarily occurs through the binding pathway of the R2 and R2* conformations. The R1/R2 equilibrium rapidly shifts towards R2. The intermediate R1* is also formed, but its population remains negligible due to its fast rearrangement rate (ke)which is significantly faster than its formation rate (kc).

These results suggest two aptamer conformations in equilibrium while one (R2) facilitates faster ligand binding. However, in the second binding step its intermediate R2* requires more extensive rearrangement towards the final bound conformation (B). In contrast, the conformation of R1 slows down initial interaction with the ligand, while its intermediate R1* allows for rapid conformational rearrangement.

### G12 folds correct, but requires magnesium for binding of doxycycline

In the next set of experiments, we investigated whether doxycycline binding to G12 depends on a magnesium-stabilized tertiary structure, similar to what has been shown for the tetracycline binding aptamer.^55^ To this end, we performed force-ramp measurements with the short stem construct in the presence of doxycycline at varying magnesium concentrations (Fig. S8h). In the absence of both magnesium and doxycycline, the folding pathway is essentially identical to that observed in the presence of Mg (Fig. 4C left). Melting curve analysis of the aptamer showed no dependency on Mg^2+^ for the structure of G12 to fold (Fig. S10c-d, Table S10). The lower overall unfolding forces are likely due to the reduced salinity / ionic strength without magnesium. In force-jump experiments, we found that, at 50 µM magnesium and 1 µM doxycycline, only ∼30% of aptamers were bound by doxycycline, indicating that magnesium concentration is still limiting. At 5 mM Mg, however, the binding probability increased to >80%. To further quantify the magnesium dependence of binding, we performed FTS at varying magnesium concentrations (Fig. S5g-h). In the absence of magnesium, no fluorescence was measured, indicating a lack of doxycycline binding to G12, while fluorescence increased progressively with magnesium concentration. With detectable binding emerging at 40 µM magnesium, the results confirm a dependency on magnesium for binding doxycycline.

## Discussion

We succeeded with identifying regulatory RNA aptamers by performing two parallel *in vitro* selections and subsequent *in vivo* screening. Starting with the identical library allowed us to directly compare outcome of the two different SELEX methods and gain valuable insights into their selection dynamics. Both selections enriched sequences binding to doxycycline. However, already the pre-selection step of the *in vivo* screening, enriching candidates not interfering with translation, revealed strong differences. The conventional SELEX showed a trend to enrich sequences generally interfering with translation initiation. Such effects are most likely caused by the formation of structures that are already too rigid in absence of the ligand to allow the ribosomal pre-initiation complex to pass the structure.^56^ In the subsequent screening, only a single clone of G12 showed considerable regulatory activity in the pools from the conventional SELEX, suggesting no enrichment of aptamer sequences with conformational changes upon ligand binding. This is in line with previous findings that regulating aptamers are rarely found in conventional SELEX approaches.^42^ In turn, the enrichment of C01 in round C7 of the Capture-SELEX indicates a very efficient enrichment of regulatory sequences through the addition of conformational change upon ligand binding as a selection parameter. The insights gained through NGS analysis further supported this finding with the rapid enrichment of the regulatory aptamer C01 over the selection process of the Capture-SELEX. Together with the observed enrichment of only a few motifs, it appears that Capture-SELEX effectively enriches RNA aptamers that undergo conformational changes upon ligand binding. Also, G12 would have most likely been found enriched in the Capture-SELEX if it would have been able to effectively hybridize to the capture-oligonucleotide. In contrast, none of the regulatory aptamers showed efficient enrichment in the conventional SELEX. Especially C01 showed fluctuating abundance over the course of the selection with no clear trend for enrichment. These findings suggest an unfavourable selection pressure in the conventional SELEX for the regulatory aptamers that effectively bound doxycycline but have been overcome by other sequences simply binding doxycycline. The overall less stringent selection dynamic becomes also apparent in the overall more diverse motif composition found in the rounds of the conventional SELEX. With more motifs identified and many motifs with considerable abundance in the last rounds of selection it appears that the conventional SELEX creates a lower selection stringency, even when introducing pre-selection, counter-selection and reduced ligand concentration, than Capture-SELEX. Overall, these results demonstrate a more efficient enrichment of regulatory aptamers through Capture-SELEX, considerably increasing the chance of their identification by subsequent screening. With these results, we hereby provide clear proof for the effectiveness of RNA Capture-SELEX for the selection of regulatory RNA aptamers suited for riboswitch engineering, as previously postulated.^34^

With G12 we obtained an aptamer with outstanding regulatory performance. Its high affinity of around 4 nM to doxycycline and complete discrimination of tetracycline are optimal prerequisites for efficient gene regulation. Its high ligand affinity is highlighted by a very low off-rate koff of approx. 0.0055 s^-1^ (mean binding duration of 182 s) and an on-rate kon of approx. 0.42 s^-1^ µM^-1^ measured with single-molecule force spectroscopy. As predicted by folding simulations and confirmed by mutational analysis and chemical probing, the aptamer folds into a T-shape secondary structure with a central three-way junction and a twelve and six nucleotide loop at each end. Correct folding of the aptamer, in contrast to the tetracycline aptamer,^55^ is independent of magnesium ions while it requires magnesium to bind doxycycline. While the upper part of the aptamer (spanning from L2 to L3) is mostly mutation intolerant, the P1 can be freely exchanged. This allowed for easy application of G12 for gene regulation in yeast and human cells. The comparative measurements in yeast clearly showed the outstanding regulatory performance of G12, by far outcompeting the other regulatory aptamers. Likewise, in human cells, G12 outperformed the tetracycline aptamer with an earlier reaction to its ligand and an overall higher dynamic range, while it was tested without further adaptations in a riboswitch context that was optimized for the tetracycline aptamer.

The aptamer G12 bases its outstanding regulatory properties on a novel and unique structural dynamic, stabilising an intermediate structure upon binding doxycycline autonomous from its P1 stem. Single-molecule force spectroscopy was performed to unravel this novel dynamic. As shown by force-ramp experiments, G12 does not show stabilisation of its P1 stem upon ligand binding, as it was observed for the tetracycline aptamer and other previously tested aptamer domains.^57,58^ The stabilisation of the P1 stem is a frequent observed reaction to ligand binding in regulatory aptamers and represents their mechanistic basis of regulation. This observation applies both to synthetic and natural aptamer domains.^59,60^ For the aptamer G12, instead a remarkable stabilisation of an intermediate structure (BI1) within the aptamer occurs. Although this stabilised structure has an overall stabilising effect on the aptamer, the intermediate structure possesses substantial stability even after unfolding of the P1 stem. This overall stabilisation of the aptamer structure is supported by the stabilising effects revealed by SHAPE-MaP in the presence of doxycycline. With the stabilisation observed in a construct of the aptamer without a P1 stem, this proves that G12 binds doxycycline and stabilises its structure completely independent from its P1 stem.

Results of the constant-force experiments showed that the stabilised intermediated structure (BI1) only appears after previous folding of the intermediate state 1 (I1), strongly suggesting conformational capture as the mode of binding. The monitored transitions between the different conformational states of G12 draw a clearly structured folding pattern. From an unfolded state, the aptamer either misfolds or forms the intermediate structure 2 (I2) first. Subsequently, the aptamer forms the intermediate structure 1 (I1) and then folds into its completely folded conformation (Fol). Only the folded intermediate 1 (I1) or the fully folded aptamer (Fol) are able to bind doxycycline and possess the increased stability as it was observed in the force-ramp experiments. Based on the mutational analysis and chemical probing, this binding competent structure is formed between the loop L2 and L3. With the lowest mutation tolerance, unexpected low nucleotide reactivity and highest stabilisation upon ligand binding, the L2 and L3 loop appear to be directly engaged in binding doxycycline. Furthermore, as shown by the contour length calculation, the stabilised intermediate structure (BI1) has a length of around 28 nucleotides. This would fit a section of the aptamer between loop L2 and L3 including the most unreactive and stabilised positions that are likely to form the binding motif. A complex binding pocket formed by this section while fully enclosing doxycycline, would explain the observed fluorescence recovery upon binding, the remarkable stabilisation of the structure and binding competence independent of the P1 stem. Overall, the results suggest a cooperative binding between the loops L2 and L3 as the most likely scenario for interaction with doxycycline, resulting in this complex representing the observed stabilised intermediate structure BI1.

When binding doxycycline, G12 undergoes a two-step binding dynamic that can take place in two ways as shown by the stopped flow spectroscopy. Similar behaviour of ligand binding dynamics has been described for other regulatory aptamers,^35–37^ but none of these aptamers possessed two binding pathways starting from two unbound conformations as observed for G12. Together with a fast first binding step, this leads to a binding pathway through R2 - R2* being the main binding dynamic. In comparison of the two binding pathways, it becomes apparent, that R1 has the slower initial binding step, but a substantial faster second binding step, compared to R2. This indicates a greater conformational rearrangement in R1 during the initial binding step, whereas R2 undergoes greater rearrangement in the second binding step. In general, the stopped flow spectroscopy results depict with G12 an aptamer that possess two binding competent conformations that both lead to a two-step binding with considerable conformational rearrangement and a bound conformation that shows minimal release of the ligand.

When comparing the difference in conformational rearrangement upon doxycycline binding of R1 and R2, it seems reasonable to assume that their conformational dynamic during binding must be detectable in the structure stability observed with single-molecule force spectroscopy. In these results two conformations are binding competent, the intermediate 1 and the fully folded aptamer. When presuming that the final bound conformation (B) is the final state of binding, all binding dynamics would have to undergo rearrangement to reach this conformation. Starting from the intermediate conformation 1 (I1), as the proposed minimal binding competent conformation, the aptamer would have to undergo further folding to reach a completely folded state which would require considerable structural rearrangement. In contrast, the fully folded conformation (Fol) would have to undergo no or little rearrangement to reach the final bound conformation (B). In comparison with the two binding pathways, it seems that R1 might reflect the binding dynamic of the fully folded aptamer. Here, the first binding step to the ligand might be hindered by the already complex tertiary structure of the aptamer, but after binding only little rearrangement is needed to tightly bind doxycycline as observed in the dynamic of the intermediate R1* to the fully bound state (B). In turn, the binding path through R2 might represent the binding starting from the minimal binding competent conformation (I1), where the binding pocket is more accessible for doxycycline, but the aptamer undergoes greater rearrangement to reach the fully bound state (B). These results provide detailed insights into the dynamics of a regulatory aptamer, connecting observations from structural stability and binding dynamics providing an explanation for the complex characteristics of aptamers necessary for their application for gene regulatory purposes.

Although RNA motifs binding doxycycline have been reported previously,^61,62^ none of these sequences has shown regulatory potential *in vivo*. Furthermore, G12 possesses the highest regulatory potential observed for a regulatory aptamer. This is based of its unique stabilisation of an intermediate conformation independent from its P1 stem. With the comprehensive characterisation of this aptamer, we provide an outstanding regulatory aptamer ready for the engineering of synthetic riboswitches that is responsive to one of the most used inducer molecules for eukaryotic cell culture. We are convinced that G12 has brought us a step closer to understand what it takes to enable an RNA molecule for regulation and that G12 will find broad and successful application in synthetic biology, biomedicine and beyond.

## Acknowledgements

The authors thank Britta Schreiber for excellent technical support with various experiments.

## Author contribution

J.H. and B.S. conceived the study and supervised the research. J.H., M.S. performed *in vitro* selections and *in vivo* screening. J.H., M.S., L.B. and T.W. performed mutational studies. J.H., A.G.J. and T.W. designed and performed NGS analysis of the selection dynamics. A.W. and M.R. designed single-molecule force spectroscopy experiments. A.W. and L.R. performed and analysed single-molecule force spectroscopy experiments. J.H. and V.G. performed and analysed SHAPE-MaP experiments. J.H. and M.S. performed and analysed ITC and fluorescence titration spectroscopy experiments. J.W. and K.S. designed stopped flow and circular dichroism experiments. K.S. performed and analysed stopped flow and circular dichroism experiments. J.H. and L.B. tested riboswitch performance in yeast. R.W.B. and O.B. tested riboswitch performance in human cells. J.H. wrote the manuscript with input from A.W., M.R., A.G.J., K.S. and B.S..

## Methods

### Material

Chemicals were purchased from Carl Roth (Karlsruhe, Germany) and used without any further purification unless otherwise stated. Molecular biology enzymes were purchased from New England Biolabs (Ipswich, USA). Oligonucleotides were either synthesized by Sigma-Aldrich (St. Louis, USA) or Microsynth (Balgach, Switzerland). Single RNA and DNA nucleotides were purchased from Sigma-Aldrich (St. Louis, USA). ^32^P-α-ATP was purchased from Hartmann Analytic (Braunschweig, Germany). Epoxy-activated Sepharose 6B was purchased from Cytiva (Marlborough, USA). Poly-Prep Chromatography columns were purchased from BIO-RAD (Hercules, USA). T7 RNA polymerase and Taq DNA polymerase are homebrew.

### Pool preparation

DNA template for the Capture-SELEX pool was assembled by PCR with 2 µM of the primers Pool_fw and Pool_rev and 70 nM of the “reverse template” (Table S11) in 20 mM TRIS pH 8,8, 10 mM (NH4)2SO4, 10 mM KCl, 2 mM MgSO4 and 0,1 % (v/v) Triton X-100 using Taq polymerase (homebrew). The PCR amplification was carried out with an initial denaturation of 2 minutes at 96°C, followed by 5 cycles of 30 seconds at 96°C, 30 seconds at 57°C and 30 seconds at 72°C with a final 3 minutes at 72°C. A T7 promoter upstream of the pool was used for *in vitro* transcription by T7 RNA polymerase (homebrew). To use the identical RNA pool for both selections, the RNA pool was transcribed in 200 mM TRIS pH 8, 20 mM Mg(OAc)2, 20 mM DTT, 4 mM ATP / UTP / GTP / CTP, 2 mM spermidine. Transcription products were ethanol precipitated, gel purified and stored for later use.

### *In vitro* selection

After starting both selections with the same prepared RNA pool, pools were transcribed from round 2 on in 100 µL with the addition of 33 nM ^32^P-γ-ATP in 200 mM TRIS pH 8, 20 mM Mg(OAc)2, 20 mM DTT, 4 mM ATP / UTP / GTP / CTP, 2 mM spermidine and 15% DMSO using T7 RNA polymerase (homebrew), resulting in ^32^P body labelling of the transcribed RNA. Transcribed RNA was precipitated using ethanol and NH4OAc and resolved in ddH2O. Radioactivity was measured using a liquid scintillation analyser (TriCard 2800 TR, PerkinElmer, Waltham, USA) determining RNA concentration.

For column-based SELEX, epoxy-activated Sepharose 6B (Cytiva, Marlborough, USA) was used. Doxycycline was coupled according to the manufacturer’s instructions at pH 12, utilizing the hydroxy groups for immobilisation. Following the same steps but leaving out doxycycline, uncoupled column material was prepared for pre-selection. The first round of selection was performed with a non-radioactive labelled RNA pool of 1.2 x 10^16^ molecules. Every round the RNA pool first underwent pre-selection by allowing the RNA pool in 500 µL 1x Capture-SELEX buffer (CSB, 40 mM HEPES (pH 7.5), 250 mM KCl, 20 mM NaCl, 5 mM MgCl2, 0,01 Tween-20) to flow through the pre-selection column of 0.5 mL Sepharose 6B without an immobilized target, to remove any sequences binding to the column material. The pre-selection column was then washed with 1 mL 1x CSB and the resulting 1.5 mL pre-selected RNA in 1x CSB was collected and allowed to flow through 1.5 mL of doxycycline-coupled Sepharose 6B. The column was then washed either 12x or 18x with 1.5 mL 1x CSB (detailed selection parameters see Supplementary Table S1). In rounds with a counter-selection step, 4.5 mL of 500 µM kanamycin A in 1x CSB were washed through the column after 12 washing steps and followed by 6 further washing steps. RNA bound to the column was eluted by allowing 4.5 mL 500 µM doxycycline in 1x CSB to flow through the column. The eluted RNA was recovered through ethanol precipitation and resuspended in 50 µL MQ H2O. The RNA was then reverse transcribed using SuperScript II reverse transcriptase (Thermo Fisher Scientific, Waltham, USA) and amplified using Taq DNA polymerase (homebrew) with 1x ThermoPol buffer, 0.2x SSII first strand buffer, 2 mM DTT, 1.5 mM MgCl2, 0.3 mM dATP/dTTP/dGTP/dCTP and 3 µM Pool_fw and Pool_rev. The cDNA pool was then stored and used for transcription of RNA for the next round of selection. The Capture-SELEX was started with 6 x 10^12^ molecules of the prepared RNA pool and performed as previously described.^41^

### In vivo screening in Saccharomyces cerevisiae

Translation control was measured in *Saccharomyces cerevisiae* RS453α (MATα *ade*2-1 *trp*1-1 *can*1-100 *leu*2-3 *his*3-1 *ura*3-52). RNA pools from SELEX were cloned into the pCBB06 backbone via homologous recombination as previously described.^44^ The *in vivo* screening system utilized a plasmid with containing a GFP gene as readout for regulatory activity, containing the aptamer candidate in its 5’UTR, and mCherry as a constitutive expressed control. For *in vivo* screening of pools form SELEX, transformed cells were first cultured in 250 mL SCD-Ura medium (0.2% YNB w/o AA (Difco), 0.55% ammonium sulphate, 2% glucose, 12 µg/mL adenine (Sigma-Aldrich), 1x MEM amino acids (Sigma-Aldrich) in a baffled flask for 48 h at 30°C and 125 rpm shaking. 25 mL of this culture were transferred to fresh 225 mL SCD-Ura medium and cultured for another 24 h at 30°C and 125 rpm shaking to reduce the risk of multiple plasmids in a single cell. The cells were sorted using fluorescence-activated cell sorting (Beckman Coulter, Brea, USA) for viability, correct mCherry expression and GFP expression. Selected cells were transferred to SCD-Ura plates (SCD-Ura medium with 2% agarose) and cultured. Single clones from these plates were used to inoculate 200 µL per well in 96 deep-well round bottom plates (Thermo Fisher Scientific) and incubated for 24 h at 30°C and 1200 rpm shaking. 20 µL of this culture were used each to inoculate 180 µL of fresh SCD-Ura in one plate without and one with 100 µM doxycycline and incubated for another 24 h. 20 µL of the culture was transferred to 180 µL PBS for GFP and mCherry fluorescence measurements using a CytoFLEX S (Beckman Coulter, Brea, USA) as previously reported.^44^

### Mutational Studies

For all mutational studies, the plasmids were used as for the *in vivo* screening in yeast. Rational designed mutants of G12 were ordered as oligonucleotides (Sigma-Aldrich, Supplementary Table S11) and hybridized by heating to 95°C and slowly cooling down. pCBB06 was digested with NheI and AgeI and the hybridized oligonucleotides were ligated into the backbone using T4 DNA ligase. The resulting plasmids were transformed into *S. cerevisiae* RS453α via electroporation using a MicroPulser Electroporator (Bio-Rad, Hercules, USA). The cells were spread on SCD-Ura plates and cultured for 48 h at 30°C. Single clones were picked to inoculate individual wells of a 96 deep well plate containing 200 µL of SCD-Ura and cultured for 48 h at 30°C and 1200 rpm. From the pre-culture, 20 µL were used to inoculate one well each on one plate containing 180 µL SCD-Ura and on one plate containing 180 µL SCD-Ura with 100 µM doxycycline and cultured for 24 h at 30°C and 1200 rpm. For fluorescence measurements, 20 µL from each well were transferred to 180 µL PBS in a 96 well plate and measured on a CytoFLEX S (Beckman Coulter, Brea, USA) as previously reported.^44^ Measurements of individual sequences were always performed twice in biological quadruplicates.

For randomized mutation studies on the variant M10, an oligonucleotide with a mutation rate of 4,5 % per nucleotide (e.g. a position originally encoding an “A” would be synthesized as 95,5% A, 1,5% T, 1,5% G, 1,5% C) from G7 to C51 (Supplementary Table S11) was ordered from Microsynth and assembled by PCR using Q5 DNA polymerase with the primers Yeast_HR_fw and Yeast-HR_rev (Sigma-Aldrich) adding overhangs for homologous recombination into pCBB06. 1 µg of NheI and AgeI digested pCBB06 with 3-times molar excess of insert was transformed by electroporation into *S. cerevisiae* RS453α using a MicroPulser Electroporator. The transformed cells were recovered in 8 mL Sorbitol-YEPD (1% yeast extract, 2% peptone, 4% glucose, 0.5 M sorbitol) and incubated 1 h at 30°C shaking at 130 rpm. The pre-cultured cells were transferred to 100 mL SCD-Ura and incubated in a baffled flask for 48 h at 30°C and shaking at 125 rpm. 20 mL of the culture were transferred to fresh 80 mL SCD-Ura medium and cultured for further 24 h. For FACS, the original construct M10 was first measured in the absence of doxycycline to set a gate covering the measured events 33% below and 33% above the medium GFP intensity. Afterwards, the cells of the partly mutated pool were measured until 100 000 cells were sorted showing GFP expression in the set reference gate. The selected cells were transferred to 100 mL SCD-Ura and regrown for 48 h at 30°C and 125 rpm. 1 mL was transferred to 50 mL SCD-Ura containing 100 µM doxycycline and incubated for 24 h at 30°C and 125 rpm shaking. For the second sorting step, again the original construct M10 was measured first in the presence of 100 µM doxycycline. Three gates were set: One gate was the gate used for the initial sorting, defined as non-regulatory since no change in expression would have occurred in these mutants. A second gate covering all events 33% below and 33% above the GFP expression of M10 with 100 µM doxycycline, defined as functional as these mutants would have shown similar expression levels to M10. A third gate that covered all events showing GFP expression lower as the second gate, defined as improved function since these mutants would show even stronger reduction in expression compared to M10. The sorted cells from the third gate were plated on SCD-Ura plates and used for flow cytometry to confirm their switching (as described for the *in vivo* screening). The cells from the first two gates were re-cultured, the plasmids isolated, the area of the aptamer PCR amplified while adding Illumina adapters and the DNA send for NGS.

### Next-generation sequencing

For Next-generation sequencing, Illumina adapter sequences were added to the RNA pool via PCR using a Q5 DNA polymerase and primers annealing in the constant regions of the pool (Figure S1a). For each round a specific forward primer was used containing an individual 6 nucleotide barcode to allow multiplexing of samples (Table S11). The samples were sequenced by GENEWIZ (Leipzig, Germany) using an Illumina NovaSeq X sequencer (Illumina, San Diego, USA) using the P5 and P7 adapter sequences. A Python-based pipeline was developed for the extraction, mutation analysis, and visualization of aptamer sequences from high-throughput sequencing data. This pipeline first extracted aptamer regions from both forward and reverse raw sequences by identifying specific primer markers and performing reverse complementation where necessary. Extracted aptamers were then subjected to global pairwise alignment against a reference aptamer sequence, followed by quality control based on length and alignment similarity. Sequences possessing below 10 reads were filtered out.

For analysis of the mutational studies, the pipeline analysed mutations by comparing aligned aptamer sequences to the reference, quantifying base substitutions and indels. Finally, it calculated and visualized mutation ratios between two samples (e.g., maintain vs. loss functions) using Log2 fold change of frequencies, generating comprehensive plots with Matplotlib that include total mutation ratios, detailed base-specific mutation ratios with significant sites highlighted, and read coverage. All mutation counts and ratios were also exported to a CSV file for further analysis.

### Identification and analysis of motifs

To identify sequence motifs enriched during the two SELEX strategies, we performed de novo motif discovery using the MEME algorithm (MEME Suite v5.5.8,). FASTA files were generated for each selection round and SELEX strategy, containing the unique sequences identified through NGS analysis. These files served as input for MEME. Motif discovery was configured to detect up to 10 enriched motifs per selection round, with motif lengths ranging from 6 to 12 nucleotides and a runtime limit of 14400 seconds. MEME output files from individual selection rounds were consolidated into a single file per SELEX strategy. Motifs were filtered to retain only those with statistically significant E-values (< 0.05). In cases where the same motif appeared in multiple rounds, only its first occurrence was retained to prevent redundancy and bias in downstream analyses. To map motifs back to the sequences, we employed the FIMO algorithm (default p-value threshold: 1e-4). Recognizing that motifs may evolve over successive selection rounds, we aligned all identified motifs using Clustal Omega and clustered them based on phylogenetic similarity. Pairwise distances were calculated using the Damerau-Levenshtein distance (via the stringdist package), and hierarchical clustering was performed using complete linkage (hclust function). Clades were defined using cutree, with a maximum pairwise distance threshold of 0.4. This clustering approach enabled the identification of distinct motif groups, which were visualized as clades across selection rounds using the ggplot2 package in stacked bar plots. Additionally, the temporal dynamics of each motif were illustrated using a combined dendrogram and bubble plot, highlighting motif abundance across the SELEX rounds and phylogenetic similarity.

### Preparation of RNA aptamers

The aptamers were transcribed from hybridized oligonucleotides (Sigma-Aldrich, Table S11) using a T7 promoter. Run-off transcription was done in 200 mM TRIS (pH 8.0), 20 mM Mg(OAc)2, 20 mM DTT, 4 mM ATP / UTP / GTP / CTP, 2 mM spermidine using T7 RNA polymerase (homebrew) and incubating for 6 h at 37°C. Mg-pyrophosphate was dissolved in the solution by dropwise adding 0.5 M EDTA (pH 8.0), the reaction products were ethanol precipitated and separated on a 10% denaturing polyacrylamide gel. The aptamer RNA was detected by ultraviolet shadowing and the respective band cut from the gel, crushed and left over night in 300 mM NaOAc solution. The supernatant was ethanol precipitated, the RNA pellet resuspended in MQ H2O, concentration measured using a NanoPhotometer N60 (Implen, Munich, Germany) and frozen for later use.

### Isothermal titration calorimetry

Gel purified aptamers and a 100 µM ligand solution were prepared in CSB. ITC measurements were performed using a MicroCal PEAQ-ITC (Malvern Instruments, XX, XX), containing the RNA in the simple cell (200 µL) and the ligand in the syringe (40 µL). After a thermal equilibration to 25°C and an initial delay of 150 s, one initial injection of 0.4 µL was carried out. Serial injections of 2 µL were performed in an interval of 120 s at a stirring speed of 750 rpm. The thermal differences occurring between the sample and reference cell were integrated and plotted against the molar ratio of RNA and ligand. The dissociation constant (KD) from each experiment was calculated using a curve fit model provided by the MicroCal PEAQ-ITC Analysis Software (v1.1.0.1262). Each measurement was performed at least twice, and final KD values were calculated from the average of the individual experiments.

### Fluorescence titration spectroscopy

Binding-induced fluorescence of doxycycline was measured using a Fluorolog FL3-22 fluorometer (Horiba, Kyoto, Japan). Measurements were performed in 2 mL FTS buffer (20 mM KPO4 pH 7.5, 100 mM NaCl 10 mM MgCl2) with 0.5 nM doxycycline or 1 nM tetracycline and varying concentrations of aptamer. The excitation wavelength was set to 470 nm and the fluorescence spectra acquired from 450 to 600 nm in 1 nm increments with an integration time of 0.1 s and slits set to 4 nm. Measurements were taken from buffer only, doxycycline in buffer and increasing concentrations of aptamer. The aptamer was titrated in small volumes to not exceed a total volume of 4%. After each titration step, the solution was stirred and allowed to equilibrate for 5 min before data collection. The fluorescence peak area around 490 nm was averaged, normalized for background fluorescence and plotted against the RNA concentration. The dissociation constant (KD) and the maximum number of binding sites were calculated as previously described^49^ and dissociation constants for each construct calculated from two measurements.

### Control of mRNA processing in human cells

Regulation of mRNA processing in human cells was tested using a dual reporter system in HeLa cells. A total of 1.5×10^4^ HeLa cells were seeded in each well of a 96-well plate and subsequently transfected with 50 fmol of the pWHE237 vector and 50 fmol of the pRL-SV40 vector using Lipofectamine 3000 (Thermo Fisher Scientific). The vector pWHE237 was containing the firefly luciferase with the riboswitch at the 3# end of the synthetic exon, while the vector pRL-SV40 was constitutively expressing a *Renilla* luciferase for normalisation, as previously described.^23^ Four hours post transfection, DMEM with or without ligand was added and the cells were incubated for 24 h. Luciferase activity was measured with the Dual-Glo® Luciferase assay (Promega, Madison, USA). 100 µL of the Dual-Glo® reagent was added to each well and incubated for 20 min at room temperature and shaking at 450 rpm. Luminescence was measured using an Infinite M200 plate reader (Tecan, Männedorf, Switzerland). Each construct was tested three times in triplicates, and the relative light units (RLU) of the firefly luciferase were normalised using the RLU of the *Renilla* luciferase. All samples were normalized to a positive control (pWHE237 without a synthetic exon). The dynamic range was calculated as the factor of signal increase between the expression level without ligand and the respective measurement with ligand.

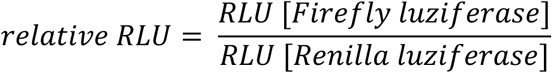

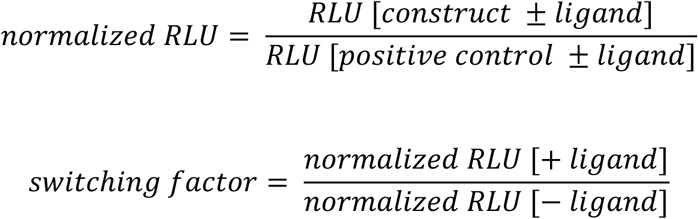

### Single-molecule force spectroscopy

#### Force Ramp Traces

For the force (F) versus extension (e) curves shown in Figure 4c, measurements were performed in so-called constant-velocity mode: the distance between the traps was increased at a fixed speed until the molecule was fully unfolded, then decreased to allow the RNA aptamer to refold. A pulling speed of 500 nm/s was used for Figure 4c, and ### nm/s for Fig.###. These traces served as molecular fingerprints, enabling identification of distinct conformational states during unfolding and refolding.

Modeling polymer elasticity in force ramp cycles: Segments of the force–extension traces corresponding to folded RNA aptamer were fitted with the extensible worm-like chain (eWLC) model.^63^

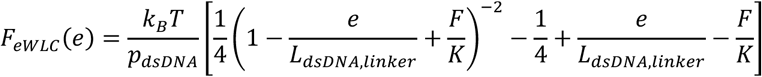

where *k*B*T* represents the thermal energy, *p*dsD*NA* is the persistence length of the dsDNA linker, *L*dsD*NA*, linker is its contour length and *K* is the elastic stretch modulus. When portions of the RNA were unfolded, the elastic response was modeled as an eWLC connected in series with a standard WLC.^64^

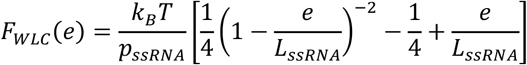

where *p*ssRNA denotes the persistence length of the unfolded ssRNA and *L*ssRNA its contour length. For fits involving unfolded ssRNA (eWLC in series with WLC), the values of *L*dsDNA,linker, *p*dsD*NA* and *K* were fixed to those obtained from the previously fitted folded state of the same force ramp trace. In addition, *p*ssRNA was fixed to 0.9 nm during fitting. The increase in unfolded contour length between successive states, determined from the WLC fits, was used to calculate the number of base pairs that had opened.

#### Passive-Mode Traces

To more accurately identify intermediate states, determine their lifetimes, and map a detailed folding/refolding pathway, we conducted passive-mode experiments in which the distance between the optical traps was held constant. This setup enabled real-time observation of RNA fluctuations between multiple intermediate states near equilibrium (Fig. 4d). In the resulting force–time traces, higher forces correspond to more folded states and lower forces to more unfolded states, as folding shortens the construct and pulls the beads out of the trap, thereby increasing the load. Each data point was assigned to a specific state using Hidden Markov Modeling (HMM).^52^ The resulting state-assigned time trajectories were used to calculate transition rates^65^ and to reconstruct the folding network (Fig. 4e).

### Off-Rates and On-Rates Determination

#### No-Stem Construct

To determine the off-rates of Doxycycline binding to the no-stem aptamer construct, we analyzed the HMM-assigned passive-mode traces. As illustrated in Figure S9a and S9e, Doxycycline-bound states are distinguished by a markedly longer lifetime on the force level of I1. Because the mean lifetimes of the unbound I1 and bound BI1 states differ by more than an order of magnitude, Doxycycline-bound events can be readily identified. The off-rate (*k_off_*) was calculated as the reciprocal of the average bound-state lifetime ⟨τ*_bound_*⟩:

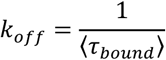

Measured off-rates (Fig. S9c) showed no significant dependence on applied force; therefore, the zero-force off-rate *k_off_*_,0_ was approximated as the mean of the force-dependent values (Table S###).

The apparent on-rate (*kon*) was obtained from the reciprocal of the average unbound-state lifetime, normalized by the Doxycycline concentration [*D*]:

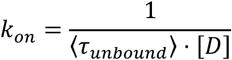

Because the binding rate to the unfolded state is significantly lower than to I1, *kon* decreases with increasing force (gray data points, Fig. S9d), where the molecule spends more time unfolded. To extract the on-rate for the native folded I1 state, the calculation was adjusted to

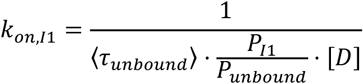

where *P*_I1_ is the probability of being in state I1 and *P_unbound_* is the probability of being unbound. This adjusted on-rate (*k_on,I1_*) showed no force dependence (Fig. S9d), and the zero-load value *k_on_* was calculated as the average of the measured *k_on_*. The dissociation constant at zero load *KD,0* was then determined as

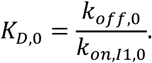

#### Short-Stem Construct

The kinetics of the short-stem construct involving the folded state were too slow to apply the passive-mode rate determination used for the no-stem construct. Instead, we performed force-jump experiments. In these experiments, the molecule was initially held at a low trap distance, producing a low force and maintaining the folded state. The trap separation was then rapidly increased. The unfolding profile revealed whether the aptamer had been bound or unbound before the jump: a short lifetime at the I1 level indicated an unbound state, whereas a long lifetime indicated a bound state (see Fig. S9e for more details).

To measure off-rates, experiments were initiated in a channel containing a Doxycycline concentration at least two orders of magnitude higher than the expected *KD*, ensuring a high probability of starting in the bound state. The construct was then rapidly transferred to a channel without Doxycycline, and after a defined waiting period, a force jump was applied to assess whether the ligand remained bound to the folded aptamer. Two waiting times were tested (∼50 s and ∼100 s). This process was repeated, and for each jump, the lifetime of the state on the I1 level was recorded. The resulting lifetime distribution exhibited a double-exponential form. Because a brief delay (∼ms) is required to reach the final trap separation and a finite observation window is imposed, lower (T1) and upper (T2) cutoffs were incorporated into the double-exponential cumulative distribution function:

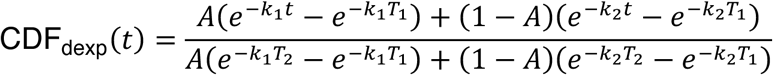

From the fit, the amplitude A was obtained, representing the fraction of bound events. Using the amplitude and the corresponding waiting time *t_wait_*, the off-rate for the folded state of the short-stem construct was calculated from a single-exponential cumulative distribution function:

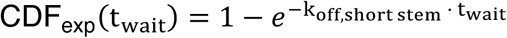

To measure the on-rates, similar jump experiments were performed. In this case, jumps were carried out in a single channel containing either 0.5 or 1 µM Doxycycline. After a defined waiting time of 4, 7 or 10 s, the force was rapidly increased from a low level to a higher level to assess whether the ligand was bound prior to the jump. Following aptamer unfolding, the force was reduced again to allow the molecule to refold into its native I1 state, enabling Doxycycline binding from the surrounding solution.

This cycle was repeated, and the dwell times on the force level of state I1 were recorded. The resulting distributions again followed a double-exponential form. The fit amplitude *A* corresponded to the fraction of jumps in which the ligand was still unbound after the waiting period. It can be assumed that the time it takes for the aptamer to refold is small compared to the waiting time, given the rapid refolding rate to state I1 even under high force (see Fig. 4d). Moreover, because the mean bound lifetime of Doxycycline to the aptamer is at least an order of magnitude longer than the waiting time, the probability of binding and subsequently unbinding within the same interval is negligible.

Using the single exponential cumulative function described above, and normalizing the rate by the ligand concentration, we determined the on-rate to the folded state of the short-stem construct (Table S###).

The dissociation constant at zero load *K_on_* for the short-stem construct was obtained by dividing the off-rate by the on-rate as described above. The reported values in Table S### are given as the mean value ± standard error of the mean determined by error propagation method.

### Stopped flow spectroscopy

The ligand capture process was monitored using a stopped-flow (SF) π*-180 device from Applied Photophysics (Leatherhead, UK) at 20°C. The following parameters were employed: excitation wavelength (λex) = 375 nm, emission wavelength (λem) > 380 nm, fluorescence detection mode at a 90° angle to the excitation source, photomultiplier tube (PMT) voltage = 450 V, acquisition of 10,000 data points (logarithmic spacing) over 50 seconds, slit width = 1 nm, and pressure hold “on.” All measurements were conducted in SFS buffer (40 mM KPi, 200 mM NaCl, 20 mM MgCl2, pH 7.5) at an RNA concentration of 4 µM, with varying ligand concentrations of 8, 16, 24, and 32 µM (2, 4, 6, and 8 equivalents (eq)). Preliminary experiments (Fig. S10a) confirmed a fully bound state under these conditions. The experimental dead time for each measurement was estimated to be approximately 150 µs. All time traces were subjected to baseline correction, averaging, smoothing (using a weighted moving average algorithm with a window size of 50 points) and normalization (to a range of 0 to 1). Kinetic modeling (KM) analysis of the normalized ligand-dependent SF data was carried out using the DynaFit4 software.^66^ A more comprehensive explanation of the data processing methodology and the various binding models employed for analysis can be found in the Supplementary Information.

## Notes

### Competing Interest Statement

Janis Hoetzel, Max Schaefer and Beatrix Suess have applied for a patent for the aptamer. The other authors declare no competing interests.

