## Supplemental Information for "A unique mechanism explaining the outstanding performance of a newly discovered doxycycline riboswitch"

### Table of content


| | |
|---|---|
| Figure S1 | p. 1 |
| Figure S2 | p. 3 |
| Figure S3 | p. 4 |
| Figure S4 | p. 6 |
| Figure S5 | p. 8 |
| Figure S6 | p. 10 |
| Figure S7 | p. 11 |
| Figure S8 | p. 12 |
| Figure S9 | p. 14 |
| Figure S10 | p. 15 |
| SUPP Method Kinetic Modelling | p. 16 |
| Table S1 | p. 17 |
| Table S2 | p. 18 |
| Table S3 | p. 19 |
| Table S4 | p. 20 |
| Table S5 | p. 21 |
| Table S6 | p. 22 |
| Table S7 | p. 23 |
| Table S8 | p. 24 |
| Table S9 | p. 25 |
| Table S10 | p. 26 |
| Table S11 | p. 27 |

### Figure S1

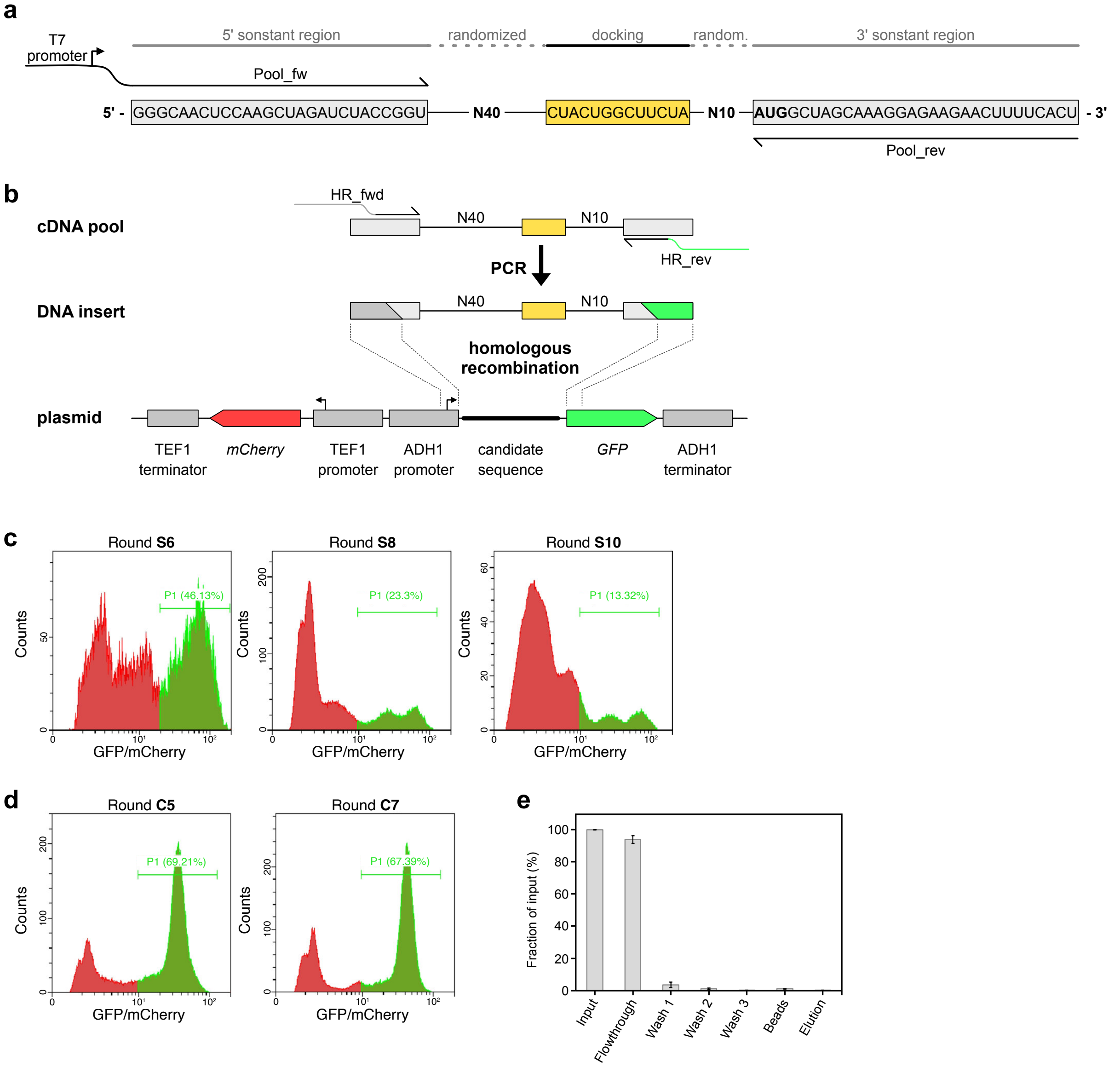


**Figure S1: Pool design & in vivo screening a**) Design of the used Capture-SELEX pool. The RNA pool consisted of a 5' and 3' constant region (grey), a N40 and N10 randomized region and the docking sequence (yellow) for hybridisation to the capture-oligonucleotide separating the randomized regions. Primer used for RT-PCR are indicted. The forward primer added the sequence of the T7 promoter to allow *in vitro* transcription of the pool for the next selection round. **b**) Schema of the procedure for the insertion of the SELEX derived pools into pCBB06 for subsequent *in vivo* screening in yeast. PCR was performed with the cDNA pool obtained from SELEX to add overhangs for the homologous recombination. The PCR product was then brought into yeast with the digested plasmid pCBB06 by electroporation. After successful homologous recombination the aptamer candidate was positioned in the 5'UTR of the GFP reporter gene. **c**) Pre-selection of aptamer candidates from rounds S6, S8 and S10 obtained from column-SELEX. Selected were viable cells with strong mCherry expression and a good GFP expression (P1, green area). The proportion of selected cells from all measured events is shown in percent. **d**) Pre-selection of aptamer candidates from rounds C5 and C7 obtained from Capture-SELEX. Selected were viable cells with strong mCherry expression and a good GFP expression (P1, green area). The proportion of selected cells from all measured events is shown in percent. **e**) Test of the aptamer G12 in the Capture-SELEX setup. Bars depict the measured radioactive signal from the $^{32}P$-labeled RNA. The candidate G12 was almost completely washed of the beads in this selection, as represented by the high fraction of RNA in the flowthrough. Remaining RNA was removed by the following washing steps, resulting in no RNA recovered in the elution step with doxycycline. This direct loss of the whole RNA in the flowthrough and washing steps indicates that the RNA has not hybridized to the capture-oligonucleotide since it was not immobilised on the beads.

### Figure S2

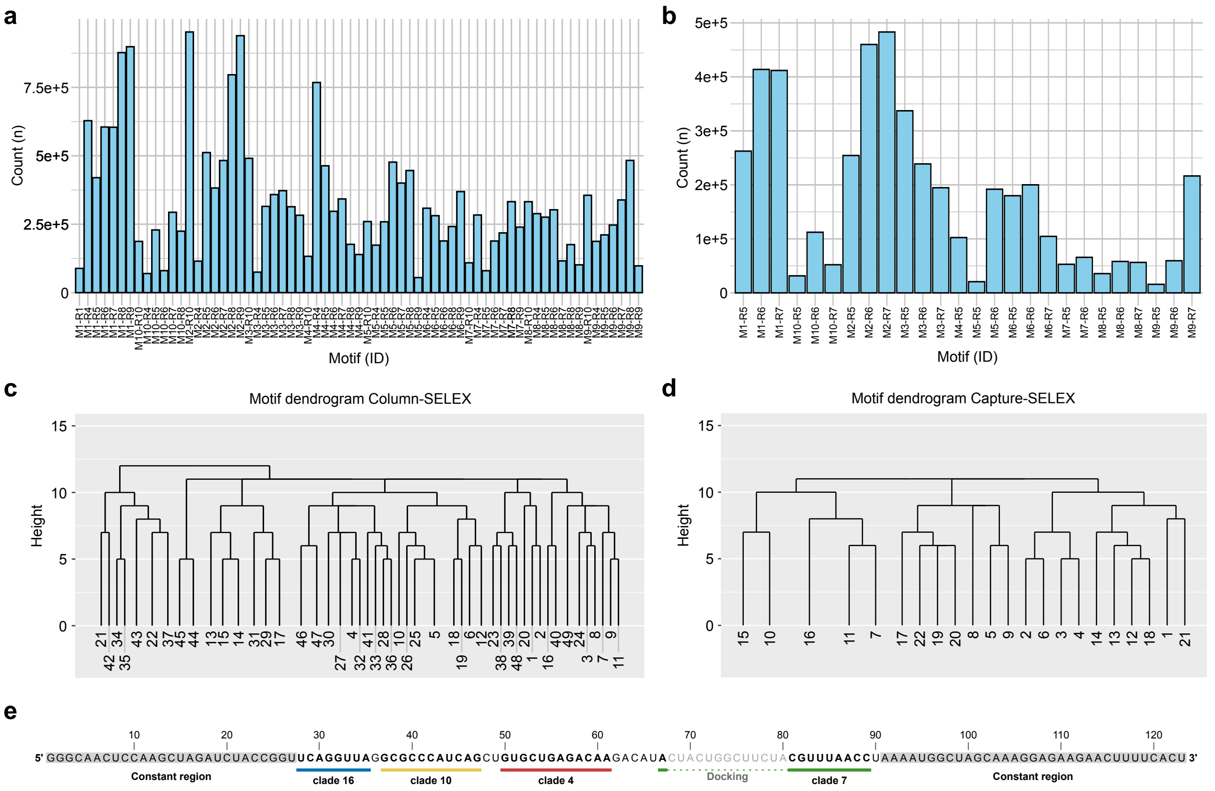


**Figure S2: Reads attributed to identified sequence motifs and their clustering to clades a**) Read counts identified for every identified motif in the column-SELEX. **b**) Read counts identified for every identified motif in the Capture-SELEX. **c**) Motif dendrogram of the motif clustering based on FIMO resulting in the motif clades 1 to 49 from the column-SELEX. **d**) Motif dendrogram of the motif clustering based on FIMO resulting in the motif clades 1 to 22 from the column-SELEX. **e**) Mapping of motif clades found in round C7 to the sequence of C01. Letters with grey background indicate the constant regions of the SELEX pool and letters in grey indicate the docking sequence.

### Figure S3

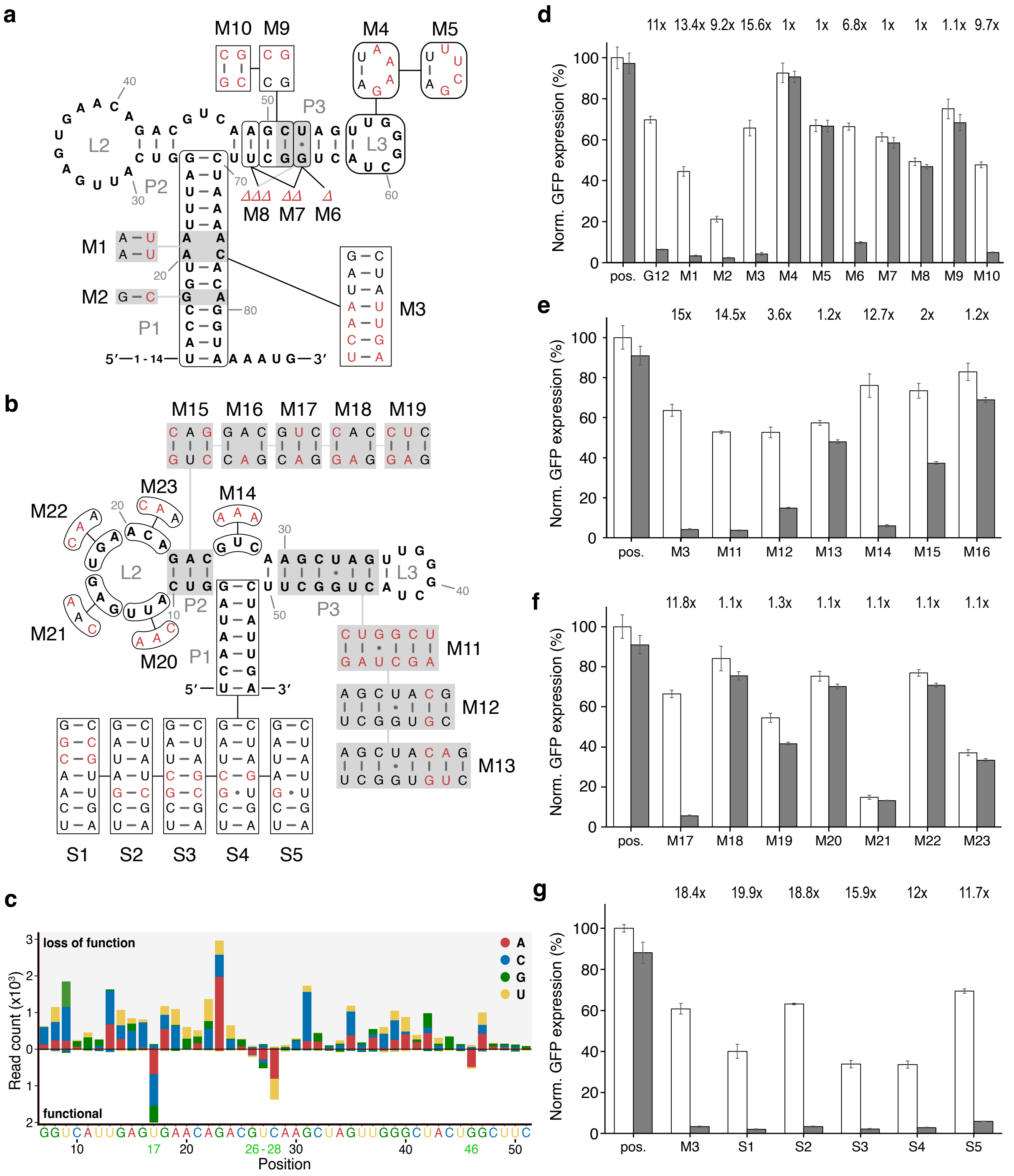


**Figure S3: Mutational studies on the aptamer G12 a**) Depiction of the predicted 2D structure (RNAfold) of G12 with the rational designed mutations tested on the aptamer. Highlighted in red are the changes introduced by the mutations. **b**) Depiction of the predicted 2D structure (RNAfold) of M3 with the rational designed mutations tested on the aptamer. Highlighted in red are the changes introduced by the mutations. **c**) Single mutant reads found in the randomized mutations screening mapped to their effect on the regulatory function and the original sequence of M3. Colours of the stacked bars indicate the changed base found at the respective position. Mutations leading to loss of regulatory behaviour are mapped as “loss of function” and mutants remaining regulatory function similar to M3 are mapped as “functional”. Positions allowing nucleotide variability without loss of regulation are highlighted by green numbers on the x-axis. **d**) GFP expression levels of G12 and its mutants M1-M10 normalized to mCherry and relative to the positive control pCBB05 without doxycycline (white bars) and with 100 $\mu$M doxycycline (grey bars). Error bars indicate standard deviation calculated from two independent measurements with biological triplicates. **e**) GFP expression levels of G3 and its mutants M11-M16 normalized to mCherry and relative to the positive control pCBB05 without doxycycline (white bars) and with 100 $\mu$M doxycycline (grey bars). Error bars indicate standard deviation calculated from two independent measurements with biological triplicates. **f**) GFP expression levels of the mutants M17-M23 normalized to mCherry and relative to the positive control pCBB05 without doxycycline (white bars) and with 100 $\mu$M doxycycline (grey bars). Error bars indicate standard deviation calculated from two independent measurements with biological triplicates. **g**) GFP expression levels of stem variants of G12 S1-S5 normalized to mCherry and relative to the positive control pCBB05 without doxycycline (white bars) and with 100 $\mu$M doxycycline (grey bars) in comparison to M3. Error bars indicate standard deviation calculated from two independent measurements with biological triplicates.

### Figure S4

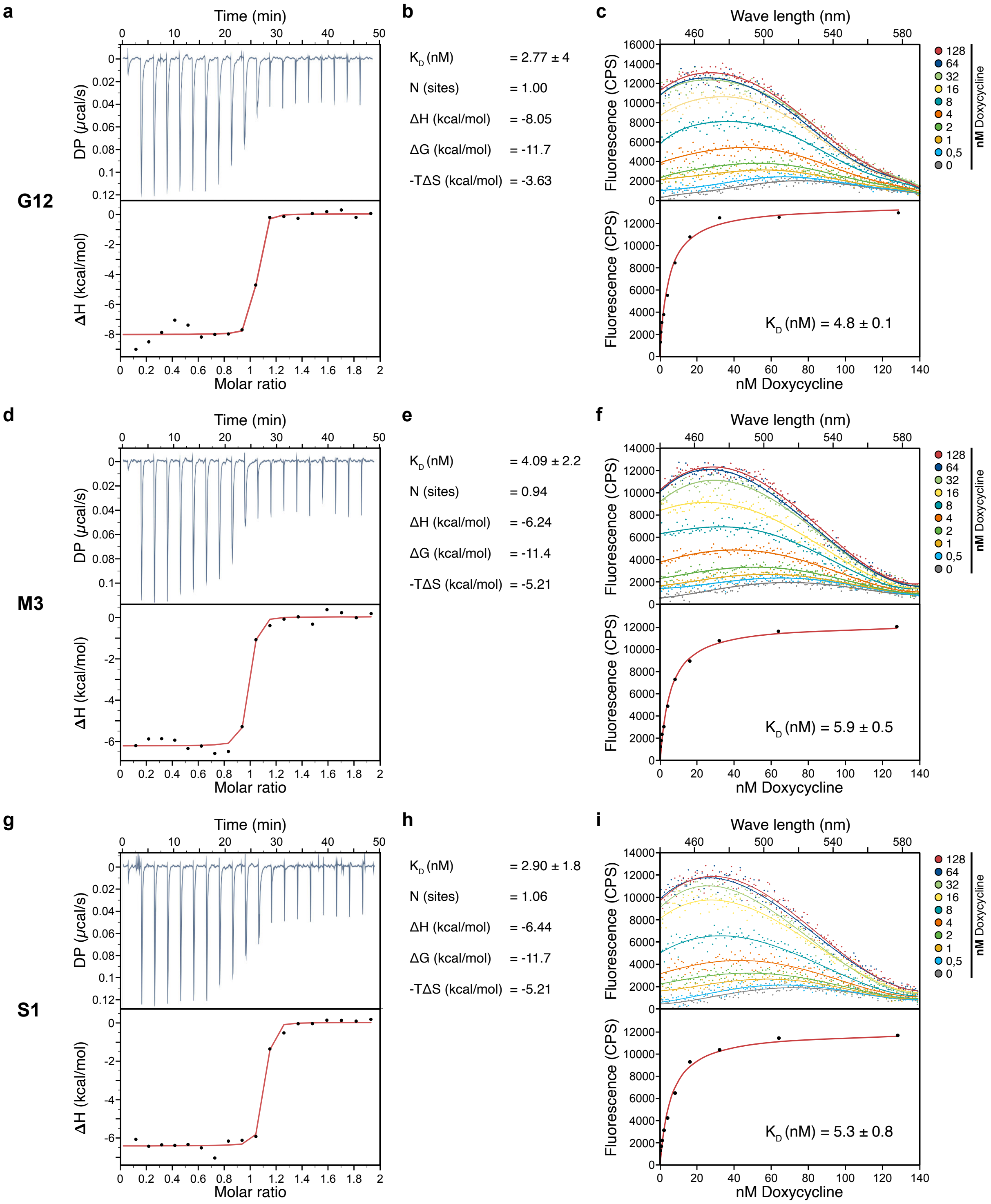


**Figure S4: $K_D$ measurements of the aptamer G12 and its variants M3 and S1 a**) Titration curve and thermogram of a representative ITC experiment measuring binding of G12 to doxycycline. Black dots depict the experimental data while the red line represents the fitted binding curve. **b**) Results of the ITC experiment measuring binding of G12 to doxycycline. **c**) Results of FTS measuring fluorescence of doxycycline when binding to G12. Shown are the measured fluorescence at different aptamer concentrations and the fluorescence levels at 490 nm wavelength used for calculation of the $K_D$. The calculated $K_D$ of the respective measurement is given below the curve. Black dots depict the experimental data while the red line represents the fitted Hill equation. **d**) Titration curve and thermogram of a representative ITC experiment measuring binding of the G12 variant M3 to doxycycline. **e**) Results of the ITC experiment measuring binding of M3 to doxycycline. **f**) Results of FTS measuring fluorescence of doxycycline when binding to M3. **d**) Titration curve and thermogram of a representative ITC experiment measuring binding of the G12 variant S1 to doxycycline. **e**) Results of the ITC experiment measuring binding of S1 to doxycycline. **f**) Results of FTS measuring fluorescence of doxycycline when binding to S1.

### Figure S5

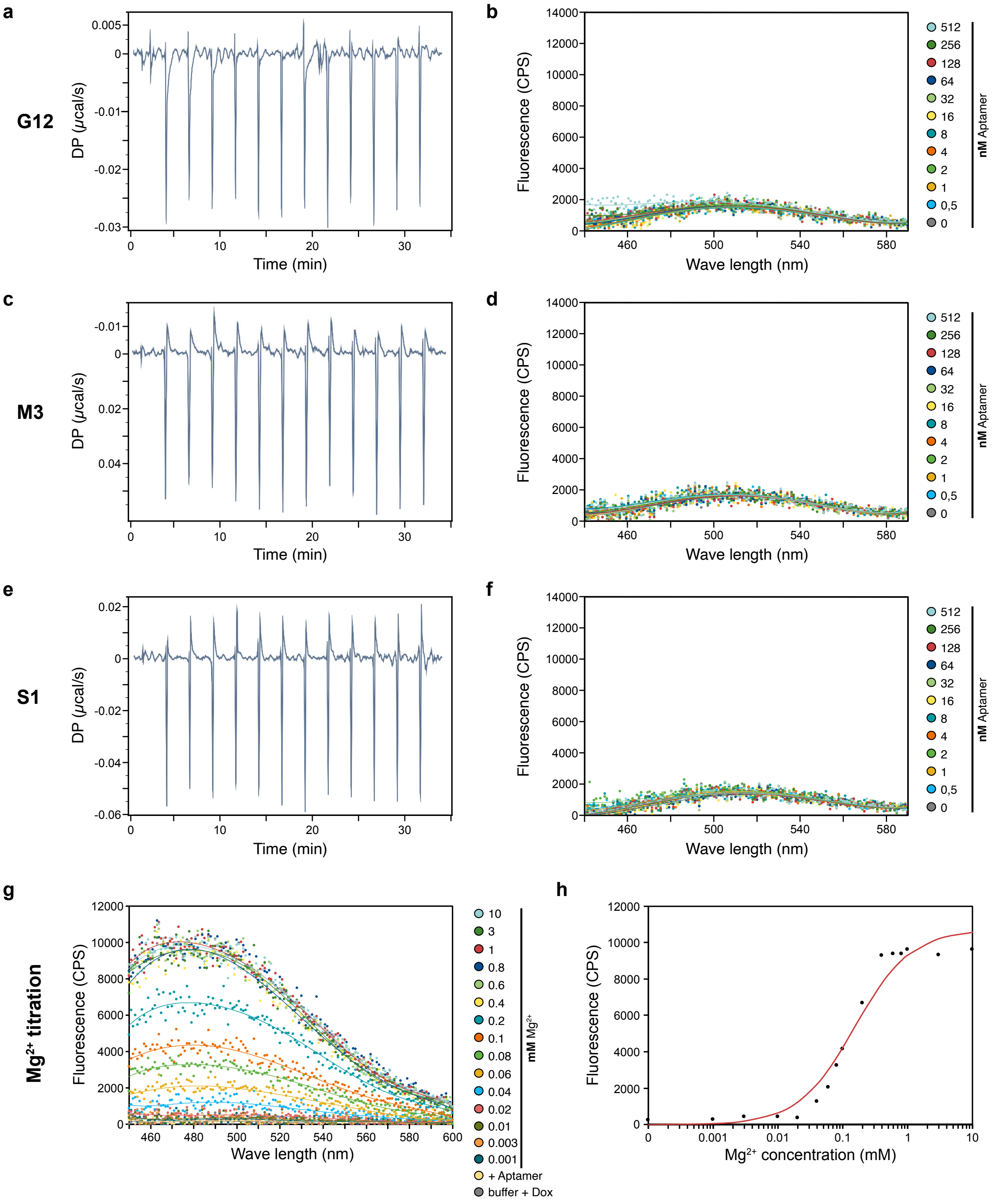


**Figure S5: Testing binding of the aptamer G12 and its variants M3 and S1 to tetracycline and testing magnesium dependence of G12 for the binding to doxycycline a**) Titration curve of ITC titrating tetracycline to G12. No signs of binding were observed. **b**) Fluorescence signal when adding G12 to tetracycline in an FTS experiment. No increase of fluorescence was observed up to an aptamer concentration of 512 nM. **c**) Titration curve of ITC titrating tetracycline to M3. No signs of binding were observed. **d**) Fluorescence signal when adding M3 to tetracycline in an FTS experiment. No increase of fluorescence was observed up to an aptamer concentration of 512 nM. **e** Titration curve of ITC titrating tetracycline to S1. No signs of binding were observed. **d**) Fluorescence signal when adding S1 to tetracycline in an FTS experiment. No increase of fluorescence was observed up to an aptamer concentration of 512 nM. **g**) Test of magnesium-dependency of G12 for binding to doxycycline. The fluorescence on the y-axis was observed for a range of magnesium concentrations from 1 $\mu$M to 10 mM. First signs of fluorescence appeared with a magnesium concentration of 40 $\mu$M. **h**) Based on the experimental data of the FTS experiment with varying magnesium concentrations, a Hill curve (red) was modelled based on the measured fluorescence (black dots) reassembling binding of doxycycline to G12 at varying magnesium concentrations.

### Figure S6

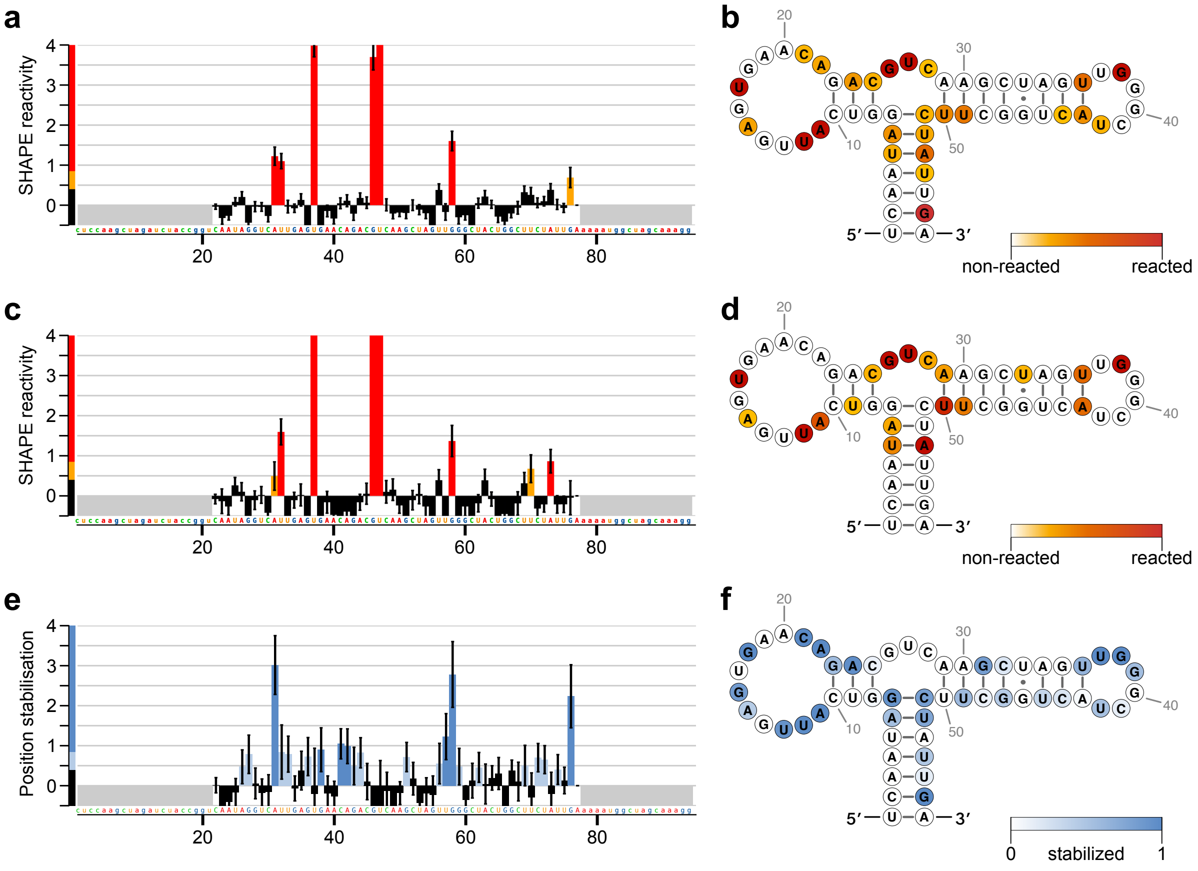


**Figure S6: Structural insights by SHAPE-MaP a**) SHAPE reactivity nucleotide positions of M3 in the absence of doxycycline. The normalized reactivity of individual nucleotide positions (x-axis) is shown on the y-axis and indicated by colour. **b**) Normalized reactivity in the absence of doxycycline mapped to the secondary structure of M3. Reactivity is indicated by colour, with the highest reactivity represented by dark red. **c**) SHAPE reactivity nucleotide positions of M3 in the presence of doxycycline. The normalized reactivity of individual nucleotide positions (x-axis) is shown on the y-axis and indicated by colour. **d**) Normalized reactivity in the presence of doxycycline mapped to the secondary structure of M3. Reactivity is indicated by colour, with the highest reactivity represented by dark red. **e**) Stabilisation of nucleotide positions of M3 calculated from the position reactivity with and without doxycycline. Stabilisation was defined as reduced reactivity in the presence of doxycycline compared to the reactivity in the absence of doxycycline. The calculated stabilisation of individual nucleotide positions (x-axis) is shown on the y-axis and indicated by colour. **f**) Calculated position stabilisation in the presence of doxycycline mapped to the secondary structure of M3. Reactivity is indicated by colour, with the strongest stabilisation indicated by dark blue.

### Figure S7

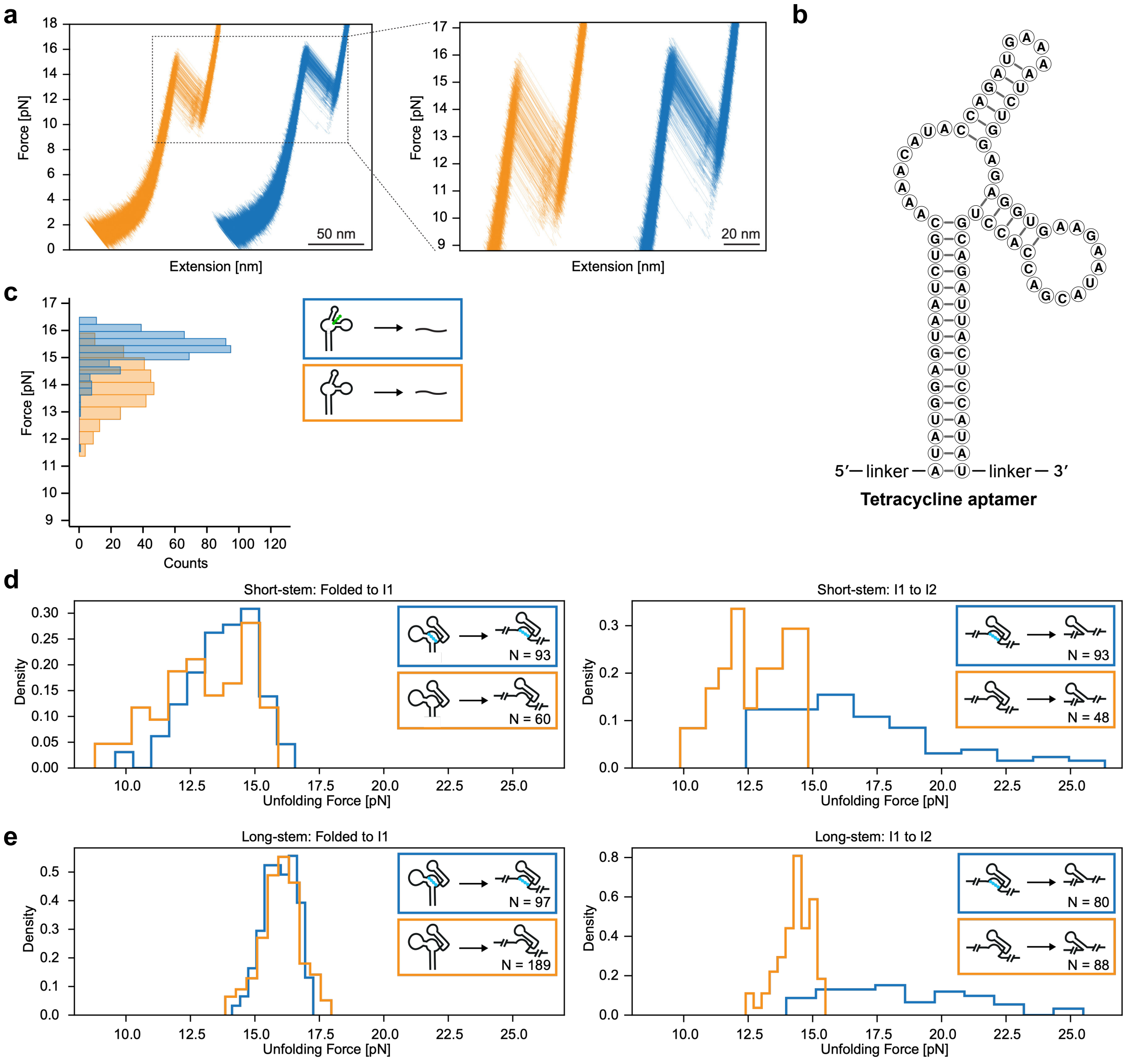


**Figure S7: Force ramp experiments of tetracycline and stabilisation of BI1 in Short-stem and Long-stem G12 constructs a**) Force-ramp unfolding traces of the tetracycline aptamer in absence (orange) or presence of tetracycline (blue). **b**) Secondary structure of the tested tetracycline aptamer. Indicated at the 5' and 3' end are the linkers used for immobilisation of the RNA that were not interfering with the folding of the aptamer. **c**) Force histogram of the unfolding of the tetracycline aptamer unbound (orange) and bound to tetracycline (blue). Clearly recognisable is the increase in force required to unfold the P1 stem. **d**) Force histogram of the unfolding of the Short-stem construct of G12 in the absence (orange) or presence of doxycycline (blue). While no difference in force required to unfold the P1 stem (Folded to I1), the unfolding of the bound intermediate 1 (I1 to I2) requires substantial more force in the presence of doxycycline. **e**) Force histogram of the unfolding of the Long-stem construct of G12 in the absence (orange) or presence of doxycycline (blue). While no difference in force required to unfold the P1 stem (Folded to I1), the unfolding of the bound intermediate 1 (I1 to I2) requires substantial more force in the presence of doxycycline.

### Figure S8

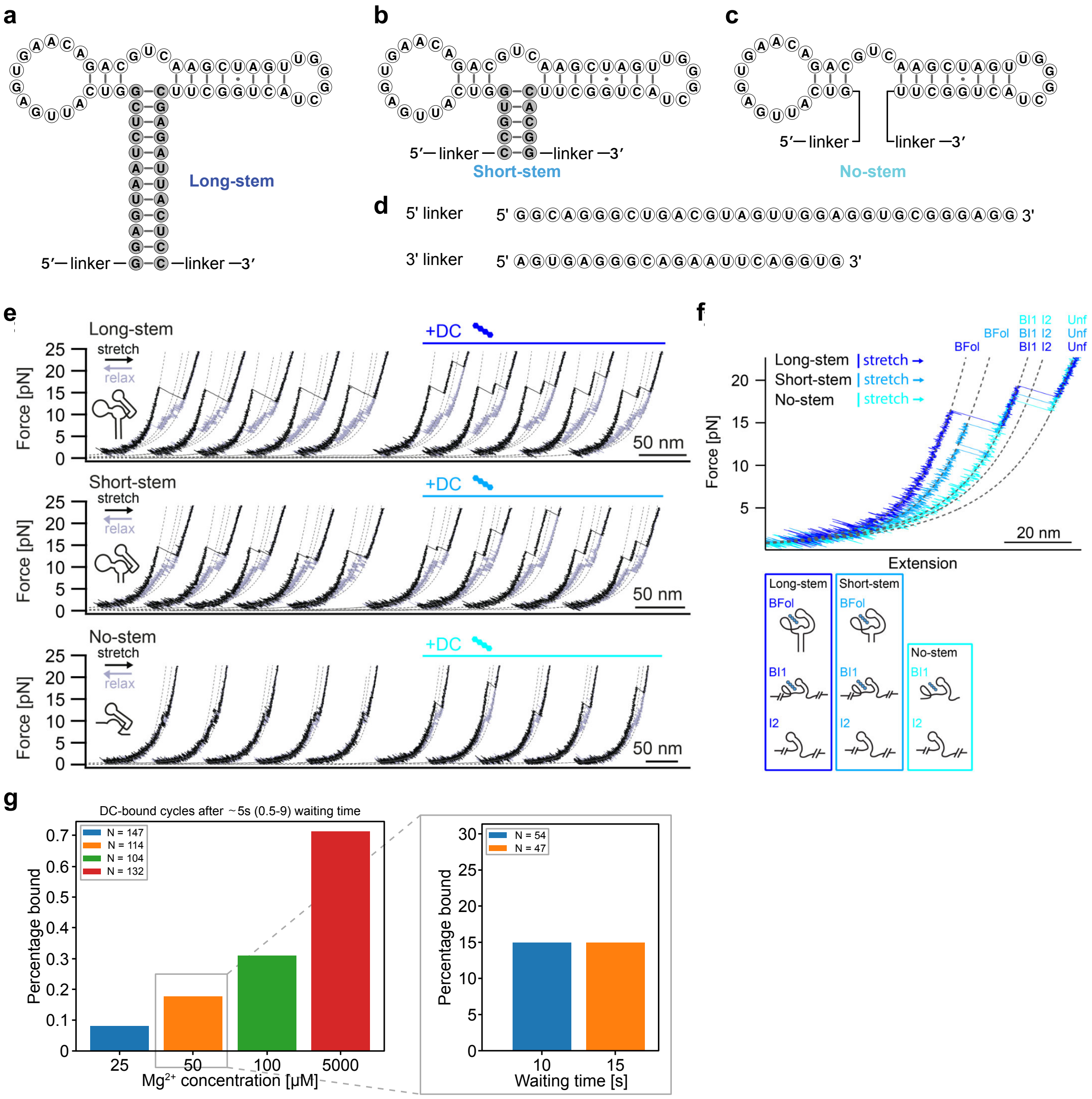


**Figure S8: Force ramp experiments of G12 constructs testing influence of P1 stability and magnesium concentration a**) Predicted secondary structure of the "Long-stem" construct of G12. The linkers necessary for immobilizing the RNA to the setup are indicated at the 5' and 3' end of the sequence. **b**) Predicted secondary structure of the "Short-stem" construct of G12. **c**) Predicted secondary structure of the "No-stem" construct of G12. **d**) Linker sequences used for the immobilisation of the RNA to the molecular tweezer setup. **e**) Representative force-ramp traces of the unfolding of the three constructs of G12 with varying P1 stems. The left five traces were measured in the absence and the left five in the presence of doxycycline. **f**) Overlay of representative force-ramp traces of the three G12 constructs in presence of doxycycline with different P1 stems. With increasing P1 stem and stability, the initial unfolding step requires a greater energy. The stabilised intermediate occurring after unfolding of the P1 stem is present in alle three constructs and stabilises independent of the P1 stem. **g**) Representative force-ramp traces of G12 in the absence of magnesium. As in the presence of magnesium all expected intermediate states of the aptamer (Fol, I1, I2, Unf) can be observed, indicating complete folding of the aptamer. **h**) Testing binding of G12 to doxycycline at varying magnesium concentration by force-ramp measurements. Binding of doxycycline was determined by observation of the stabilisation of the BI1 intermediate structure. The percentage of the events showing binding to doxycycline increased with increasing magnesium concentration.

### Figure S9

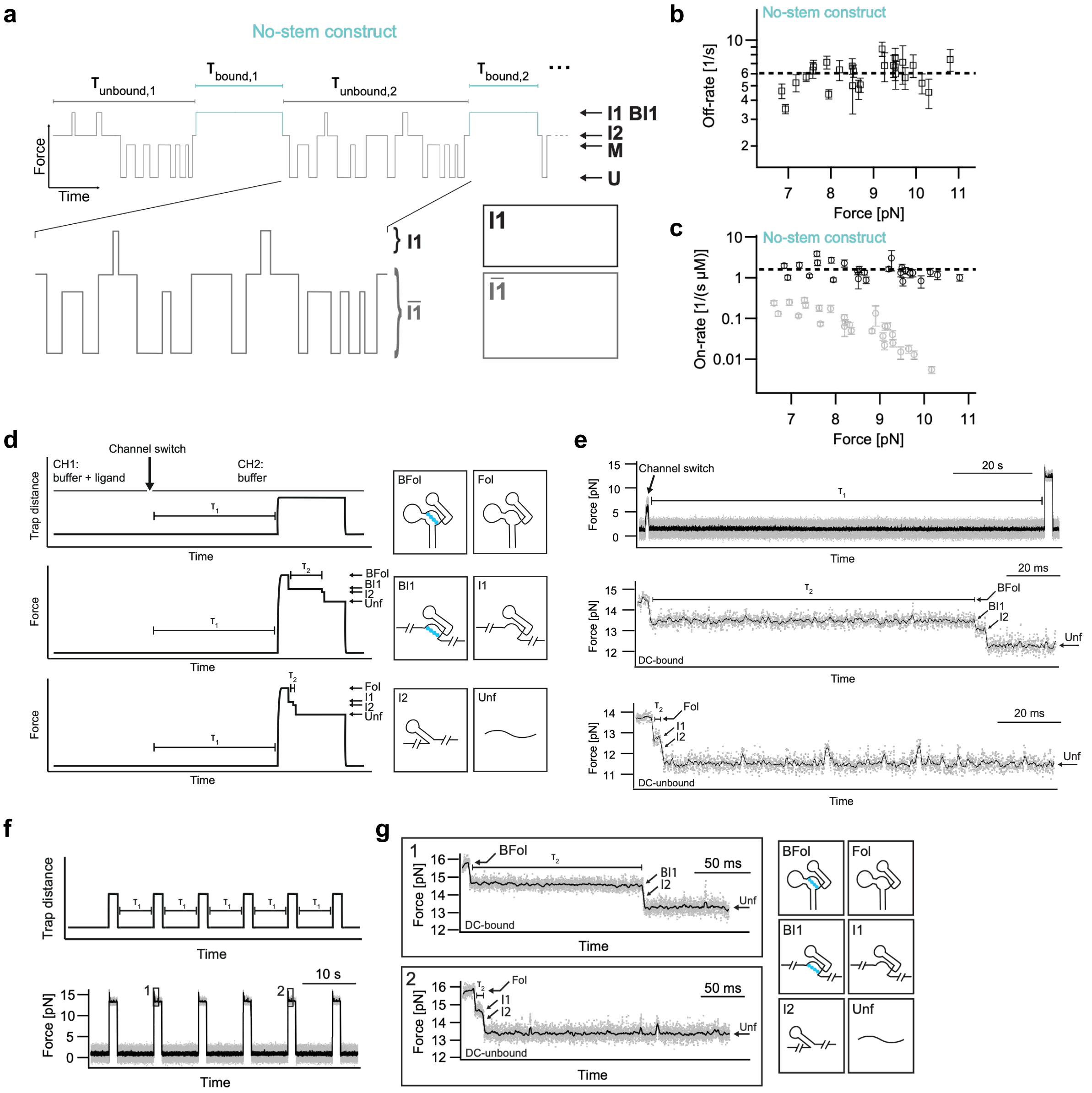


**Figure S9: Jump-force experiments for determination of ON and OFF rates of G12 a**) Schema of traces in jump-force experiments determining the OFF rate of G12 to doxycycline. After initial binding of doxycycline in microfluidic channel 1 (CH1), the aptamer bound to doxycycline is moved to channel 2 (CH2) containing no doxycycline. After a defined time ($T_1$) a force-jump is performed and the resulting resistance observed. The different intermediate strucutres of the aptamers are expected to occur depending on whether doxycycline was bound and in which conformation the aptamer was present. **b**) Representative traces of G12 bound to doxycycline (DC-bound) and unbound (DC-unbound) in the jump-force experiment. **c**) Calculated Off-rates from multiple measurements with individual molecules revealing an Off-rate of 0.0055 $s^{-1}$. **d**) Calculated On-rates from multiple measurements with individual molecules revealing an On-rate of 0.4 $s^{-1}$ $\mu M^{-1}$. **e**) Schema of traces in jump-force experiments determining the ON rate of G12 to doxycycline. The aptamer was unfolded and refolded in a microfluidic channel containing doxycycline. Force-jumps were performed after a defined time ($T_1$) and the resulting resistance of the aptamer was observed. **f**) Representative traces of G12 bound to doxycycline (DC-bound) and not bound (DC-unbound) occurring during a force-jump, making clearly distinguishable if doxycycline was bound or not.

### Figure S10

a b c d

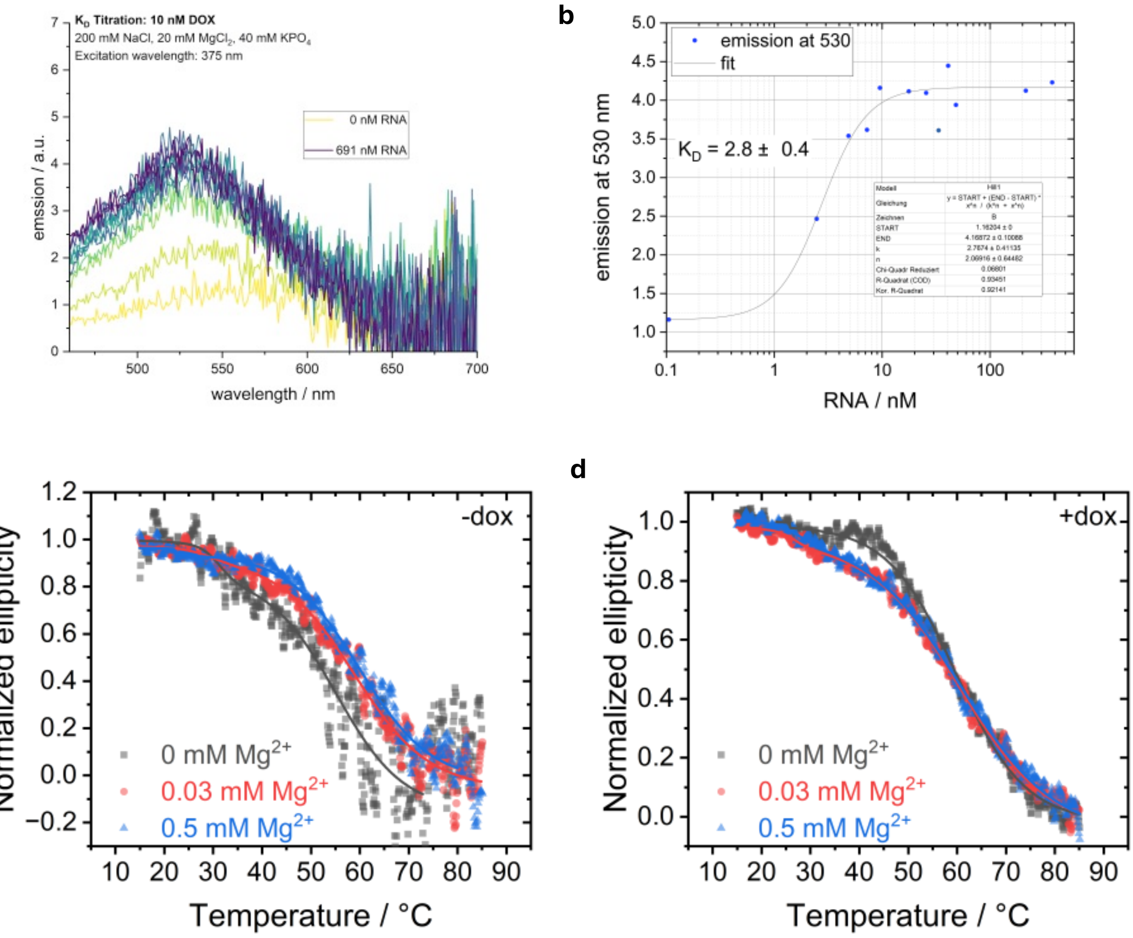


**Figure S10: Initial fluorescence titration and circular dichroism thermal melting assay of G12**
**a**) All spectra were measured using an FP-8500 fluorescence spectrometer (Jasco) in a 2x10 mm quartz cuvette, with background correction, solvent excitation correction, and reabsorption correction applied. A concentration of 10 nM doxycycline was used, and spectra were acquired at varying amounts of added G12 until saturation was reached at a constant temperature of 20 °C, with an excitation wavelength of 375 nm. The emission at 530 nm was utilized to determine the $K_D$ value using the Hill equation. A $K_D$ value of 2.8 nM was obtained. The corrected fluorescence spectra of doxycycline (yellow) and increasing amounts of G12 (color gradient towards blue). **b**) The emission at 530 nm is plotted against the added RNA concentration. The black line represents the fit via a Hill equation and yields a $K_D$-value of 2.8 nM. **c**) Circular dichroism thermal melting assay of G12 in presence of 0/0.03/0.5 mM (black/red/blue) magnesium without doxycycline and **d**) with doxycycline. The dots represent the experimental data, while the lines show the obtained fit according to equation

#### Suppl. Methods

**Kinetic modelling analysis:** The dataset was fitted with the software Dynafit4 (Kuzmič, P. (1996). Program DYNAFIT for the analysis of enzyme kinetic data: application to HIV proteinase. *Analytical biochemistry*, *237*(2), 260-273.). Dynafit4 translates symbolic reaction schemes into matrices and solves multiple equilibrium problems. The solutions of these problems can be stated as:

$$S(t) = S_0 + \sum_{i=0}^{n} r_i c_i(t) \qquad (1)$$

Here, $S(t)$ is the measured time-dependent fluorescence signal with a fixed starting offset $S_0$. $n$ is the number of distinct species involved in the reaction scheme, each characterized by a corresponding fluorescence contribution change value (response coefficient) $r$ and the concentration $c$, which varies during the binding reaction. The response coefficients are derived from the fluorescence contribution of each species. As the analyzed time traces represent the difference between the fluorescence at time t and the initial fluorescence, the response coefficient signifies the magnitude of fluorescence change associated with each species.

In this study, we employed a model selection approach incorporating the root mean square error (RMSE) as well as the Akaike Information Criterion (AIC, Burnham, K. B., and Anderson, D. R. (2002). *Model Selection and Multimodel Inference: A Practical Information-Theoretic Approach*. Springer New York). The AIC penalizes models with a large number of parameters, preventing overfitting. The AIC is calculated as:

$$\mathrm{AIC} = \left[-\log\left(\mathrm{L_p}\right) + 2\ \mathrm{p}\right] \qquad (2)$$

where p is the number of parameters in the model and *L* is the maximum likelihood. A lower AIC value indicates a better model fit. We use the □AIC-value, where the lowest AIC value is subtracted from each AIC value. Thus 0 is the smallest AIC value and the most likely model. The considered models (Table 1) represent various conformational states of the protein: $R_1$ and $R_2$ (ligand-free G12 conformations), $R_1$* and $R_2$* (ligand-associated intermediate G12 conformations), and B (ligand-bound complex).

A single binding step model proved Insufficient to accurately represent the experimental data. Therefore, we explored two-state models with starting conformations $R_1$ and $R_2$. Subsequently, an intermediate conformation was introduced to account for ligand contact.

Several binding scenarios were considered: (1) only one conformation is binding competent, binding occurs via an intermediate state, (2) both conformations are binding competent, binding occurs via the same intermediate state, and (3) both conformations are binding competent, binding occurs via separate intermediate states. Additionally, various back reactions were investigated to refine the model.

It Is Important to note that models with multiple Initial conformations can be sensitive to the initial concentration of each component. To assess the impact of initial concentrations, we explored different ratios of $R_1$ and $R_2$: 1:0, 0:1, and 1:1.

The calculated rate constants for each model are provided in the attached ZIP folder as an HTML file. The AIC and RMSE values, as determined by Dynafit4, are listed in Table S9. The models are ranked according to their ΔAIC values, with the lowest ΔAIC indicating the best fit. For each model and each individual SF time trace, we received the percentage of each population for each timepoint. In Figure 4h we show the population variation over the measuring window of all eq. of the best fitting model to illustrate the ligand binding process.

**Table S1:** Selection parameters and elution fraction of the column-SELEX

| Selection round | Washing steps | Counter-selection w/ kanamycin A ($\mu$M) | Concentration doxycycline for elution ($\mu$M) | Eluted RNA (% relative to input) |
|---|---|---|---|---|
| **S1** | 12 | 0 | 500 | - |
| **S2** | 12 | 0 | 500 | 0.01 |
| **S3** | 12 | 0 | 500 | 0.01 |
| **S4** | 12 | 0 | 500 | 0.22 |
| **S5** | 12 | 0 | 500 | 5.61 |
| **S6** | 18 | 0 | 500 | 3.98 |
| **S7** | 18 | 500 | 500 | 0.09 |
| **S8** | 18 | 500 | 500 | 1.42 |
| **S9** | 18 | 500 | 500 | 11.28 |
| **S10** | 18 | 0 | 50 | 35.03 |

**Table S2:** Selection parameters and eluted fraciton of the Capture-SELEX

| **Selection round** | Washing steps | Eluted background (% relative to input) | Concentration doxycycline for elution ($\mu$M) | Eluted RNA (% relative to input) |
|---|---|---|---|---|
| **C1** | 3 | - | 100 | - |
| **C2** | 3 | 3.07 | 100 | 2.38 |
| **C3** | 3 | 2.89 | 100 | 2.54 |
| **C4** | 3 | 3.66 | 100 | 3.05 |
| **C5** | 3 | 2.83 | 100 | 3.78 |
| **C6** | 3 | 5.35 | 100 | 18.65 |
| **C7** | 3 | 6.63 | 100 | 37.24 |

**Table S3:** Regulatory performance of G12 mutants tested in yeast

| Aptamer | Expression w/o doxycycline (%) | Expression w/ 100 $\mu$M doxycycline (%) | Fold-change |
|---|---|---|---|
| none (pCBB05) | 100 | 97.3 | 1.0 |
| G12 | 69.8 | 6.4 | 11.0 |
| M1 | 44.4 | 3.2 | 13.8 |
| M2 | 21.2 | 2.3 | 9.2 |
| M3 | 65.7 | 4.2 | 15.6 |
| M4 | 92.6 | 90.6 | 1.0 |
| M5 | 66.8 | 66.6 | 1.0 |
| M6 | 66.4 | 9.7 | 6.8 |
| M7 | 61.3 | 58.5 | 1.0 |
| M8 | 49.3 | 46.8 | 1.1 |
| M9 | 75.0 | 68.4 | 1.1 |
| M10 | 47.7 | 4.9 | 9.7 |
| M11 | 52.8 | 3.6 | 14.5 |
| M12 | 52.6 | 14.8 | 3.6 |
| M13 | 57.4 | 48.0 | 1.2 |
| M14 | 76.0 | 6.0 | 12.7 |
| M15 | 73.5 | 37.3 | 2.0 |
| M16 | 82.9 | 68.9 | 1.2 |
| M17 | 66.4 | 5.6 | 11.8 |
| M18 | 84.1 | 75.4 | 1.1 |
| M19 | 54.5 | 41.6 | 1.3 |
| M20 | 75.3 | 70.2 | 1.1 |
| M21 | 14.8 | 13.2 | 1.1 |
| M22 | 76.9 | 70.8 | 1.1 |
| M23 | 37.1 | 33.4 | 1.1 |
| S1 | 40.1 | 2.0 | 19.9 |
| S2 | 63.1 | 3.4 | 18.8 |
| S3 | 33.8 | 2.1 | 15.9 |
| S4 | 33.6 | 2.8 | 12.0 |
| S5 | 69.4 | 5.9 | 11.7 |

**Table S4:** Dose dependency of the regulatory activity of G12 and its variants in yeast

| | G12 | | M3 | | S1 | |
|---|---|---|---|---|---|---|
| *μ*M doxycycline | Expression (%) | Fold-change | Expression (%) | Fold-change | Expression (%) | Fold-change |
| 0 | 67.7 | 1.0 | 64.6 | 1.0 | 40.9 | 1.0 |
| 0.01 | 69.1 | 1.0 | 62.9 | 1.0 | 39.2 | 1.0 |
| 0.1 | 63.5 | 1.1 | 59.6 | 1.1 | 38.0 | 1.1 |
| 0.5 | 55.8 | 1.2 | 53.4 | 1.2 | 33.3 | 1.3 |
| 1 | 48.8 | 1.4 | 43.9 | 1.5 | 29.0 | 1.4 |
| 5 | 27.4 | 2.5 | 25.0 | 2.6 | 14.7 | 2.6 |
| 10 | 19.0 | 3.6 | 16.1 | 4.0 | 9.5 | 4.3 |
| 25 | 11.7 | 5.8 | 9.4 | 6.8 | 5.3 | 7.7 |
| 100 | 6.2 | 10.9 | 4.1 | 15.9 | 2.7 | 15.5 |
| 250 | 4.2 | 16.0 | 2.7 | 23.8 | 2.1 | 23.7 |
| **EC50** | **2.7 ± 0.7 *μ*M** | | **2.6 ± 0.6 *μ*M** | | **2.6 ± 0.4 *μ*M** | |

**Table S5:** Regulatory potential of different aptamers in yeast

| Aptamer | Expression w/o ligand (%) | Expression w/ 100 $\mu$M ligand (%) | Expression w/ 250 $\mu$M ligand (%) | Fold-change 100 $\mu$M ligand | Fold-change 250 $\mu$M ligand |
|---|---|---|---|---|---|
| Tetracycline “tc” | 30.7 | 8.9 | 5.5 | 3.4 | 5.6 |
| Tetracycline G6 | 31.2 | 5.1 | 4.0 | 6.1 | 7.7 |
| Tobramycin T1 | 38.8 | 4.4 | 4.0 | 8.8 | 9.6 |
| Tobramycin N1-G6 | 36.2 | 4.4 | 3.9 | 8.2 | 9.3 |
| Neomycin M4 | 27.6 | 4.8 | 3.9 | 5.7 | 7.2 |
| Doxycycline G12 | 64.3 | 6.3 | 4.6 | 10.3 | 14.1 |
| Doxycycline M3 | 61.3 | 4.4 | 3.1 | 14.1 | 20.0 |
| Doxycycline S1 | 38.2 | 2.1 | 1.5 | 18.0 | 25.0 |

**Table S6:** Contour lengths of the intermediate aptamer structures by force-ramp experiments. For all measurements the standard error of the mean (SEM) is indicated behind the values.

| Construct | Transition | Contour length (SEM) [nm] |
|---|---|---|
| Long-stem | Fol - I1 | 23.91 (0.14) |
| | BFol - BI1 | 23.69 (0.07) |
| | I1 - I2 | 6.6 (0.3) |
| | BI1 - I2 | 6.75 (0.21) |
| | I2 - Unf | 10.66 (0.18) |
| Short-stem | Fol - I1 | 14.6 (0.7) |
| | BFol - BI1 | 14.52 (0.13) |
| | I1 - I2 | 6.5 (0.3) |
| | BI1 - I2 | 6.3 (0.3) |
| | I2 - Unf | 10.82 (0.17) |
| No-stem | I1 - I2 | 6.2 (0.4) |
| | BI1 - I2 | 6.8 (0.4) |
| | I2 - Unf | 10.8 (0.3) |
| | Mis - Unf | 6.44 (0.13) |

**Table S7:** Affinities of G12 aptamer constructs binding doxycycline determined by single-molecule force spectroscopy. For all measurements the standard error of the mean (SEM) is indicated behind the values.

| Construct | On-rates (SEM) [1/(s $\mu$M)] | Off-rates (SEM) [1/s] | $K_D$ (SEM) [$\mu$M] |
|---|---|---|---|
| Short-stem G12 | 1.59 (0.15) | 6.04 (0.24) | 3.9 (0.4) |
| No-stem G12 | 0.42 (0.03) | 0.0055 (0.0003) | 0.0131 (0.0013) |

**Table S8**: A list of all 24 tested models during the comparative analysis of the ligand-binding kinetics of G12 and doxycycline. A symbolic reaction scheme as well as the starting concentration of $R_1$ and $R_2$ for each model are given.

| Model number | model scheme | $C_0$ [$R_1$] | $C_0$ [$R_2$] |
|---|---|---|---|
| 1 | $R_1 + D \rightarrow B$ | 4 | - |
| 2 | $R_1 \rightleftharpoons R_2 + D \rightarrow B$ | 2 | 2 |
| 3 | $R_2 \rightleftharpoons R_1 + D \rightarrow B$ | 2 | 2 |
| 4 | $R_1 \rightleftharpoons R_2 + D \rightarrow B$<br>$R_1 + D \rightarrow B$ | 2 | 2 |
| 5 | $R_1 + D \rightarrow R_1^* \rightarrow B$ | 4 | - |
| 6 | $R_1 + D \rightleftharpoons R_1^* \rightarrow B$ | 4 | - |
| 7 | $R_2 \rightleftharpoons R_1 + D \rightarrow R_1^* \rightarrow B$ | 4 | 0 |
| 8 | $R_2 \rightleftharpoons R_1 + D \rightarrow R_1^* \rightarrow B$ | 0 | 4 |
| 9 | $R_2 \rightleftharpoons R_1 + D \rightarrow R_1^* \rightarrow B$ | 2 | 2 |
| 10 | $R_1 \rightleftharpoons R_2 + D \rightarrow R_2^* \rightarrow B$ | 4 | 0 |
| 11 | $R_1 \rightleftharpoons R_2 + D \rightarrow R_2^* \rightarrow B$ | 0 | 4 |
| 12 | $R_1 \rightleftharpoons R_2 + D \rightarrow R_2^* \rightarrow B$ | 2 | 2 |
| 13 | $R_2 \rightleftharpoons R_1 + D \rightarrow R^* \rightarrow B$<br>$R_2 + D \rightarrow R^* \rightarrow B$ | 4 | 0 |
| 14 | $R_2 \rightleftharpoons R_1 + D \rightarrow R^* \rightarrow B$<br>$R_2 + D \rightarrow R^* \rightarrow B$ | 0 | 4 |
| 15 | $R_2 \rightleftharpoons R_1 + D \rightarrow R^* \rightarrow B$<br>$R_2 + D \rightarrow R^* \rightarrow B$ | 2 | 2 |
| 16 | $R_2 \rightleftharpoons R_1 + D \rightarrow R_1^* \rightarrow B$<br>$R_2 + D \rightarrow R_2^* \rightarrow B$ | 4 | 0 |
| 17 | $R_2 \rightleftharpoons R_1 + D \rightarrow R_1^* \rightarrow B$<br>$R_2 + D \rightarrow R_2^* \rightarrow B$ | 0 | 4 |
| 18 | $R_2 \rightleftharpoons R_1 + D \rightarrow R_1^* \rightarrow B$<br>$R_2 + D \rightarrow R_2^* \rightarrow B$ | 2 | 2 |
| 19 | $R_2 \rightleftharpoons R_1 + D \rightleftharpoons R_1^* \rightarrow B$<br>$R_2 + D \rightleftharpoons R_2^* \rightarrow B$ | 4 | 0 |
| 20 | $R_2 \rightleftharpoons R_1 + D \rightleftharpoons R_1^* \rightarrow B$<br>$R_2 + D \rightleftharpoons R_2^* \rightarrow B$ | 0 | 4 |
| 21 | $R_2 \rightleftharpoons R_1 + D \rightleftharpoons R_1^* \rightarrow B$<br>$R_2 + D \rightleftharpoons R_2^* \rightarrow B$ | 2 | 2 |
| 22 | $R_2 \rightleftharpoons R_1 + D \rightleftharpoons R_1^* \rightleftharpoons B$<br>$R_2 + D \rightleftharpoons R_2^* \rightleftharpoons B$ | 4 | 0 |
| 23 | $R_2 \rightleftharpoons R_1 + D \rightleftharpoons R_1^* \rightleftharpoons B$<br>$R_2 + D \rightleftharpoons R_2^* \rightleftharpoons B$ | 0 | 4 |
| 24 | $R_2 \rightleftharpoons R_1 + D \rightleftharpoons R_1^* \rightleftharpoons B$<br>$R_2 + D \rightleftharpoons R_2^* \rightleftharpoons B$ | 2 | 2 |

**Table S9:** Summary of the comparative analyses of the ligand-binding kinetics of G12. The obtained RMSE of the fit as well as the relative comparison via $SSQ_R$ (relative sum of squared deviations between experimental data and the theoretical fit) are given. The relative probabilities of the models are reflected by the lowest value of the likelihood criteria ΔAIC.

| Model number | $SSQ_R$ | ΔAIC | RMSE |
|---|---|---|---|
| 18 | 1.000 | 0.0 | 0.0333886 |
| 22 | 1.011 | 458.7 | 0.0335772 |
| 17 | 1.031 | 1202.0 | 0.0338940 |
| 19 | 1.031 | 1207.5 | 0.0338947 |
| 21 | 1.040 | 1564.8 | 0.0340464 |
| 16 | 1.043 | 1674.3 | 0.0340947 |
| 14 | 1.047 | 1825.0 | 0.0341607 |
| 13 | 1.047 | 1826.3 | 0.0341613 |
| 20 | 1.047 | 1858.9 | 0.0341718 |
| 15 | 1.060 | 2321.8 | 0.0343735 |
| 12 | 1.068 | 2641.7 | 0.0345121 |
| 24 | 1.083 | 3185.3 | 0.0347413 |
| 9 | 1.089 | 3396.0 | 0.0348390 |
| 11 | 1.106 | 4013.3 | 0.0351089 |
| 23 | 1.107 | 4091.7 | 0.0351372 |
| 10 | 1.108 | 4150.5 | 0.0351494 |
| 8 | 1.108 | 4107.7 | 0.0351504 |
| 6 | 1.108 | 4111.1 | 0.0351528 |
| 4 | 1.113 | 4257.9 | 0.0352173 |
| 2 | 1.122 | 4584.6 | 0.0353623 |
| 3 | 1.122 | 4584.6 | 0.0353623 |
| 7 | 1.154 | 5709.8 | 0.0358614 |
| 5 | 1.156 | 5797.5 | 0.0359026 |
| 1 | 1.232 | 8341.1 | 0.0370643 |

**Table S10:** List of determined melting points form the CD thermal melting assay

| $Mg^{2+}$ concentration | CD melting points | | | |
|---|---|---|---|---|
| | - Dox | | + Dox | |
| 0 mM | 31.3 | 55.1 | 25.8 | 59.8 |
| 0.03 mM | 29.6 | 58.8 | 26.8 | 60.4 |
| 0.5 mM | 22.1 | 59.8 | 27.8 | 60.7 |

**Table S11:** Oligonucleotide sequences

Positions labelled as "N" indicate a random distribution between all four nucleobases (A, T, G, C). Barcodes used for NGS multiplexing are indicated in **bold**. For the "G12_M3_random-template" used for the randomized mutation screening, the partly randomized section (4,5% mutation rate/nucleotide) is indicated by underlining.

| Name | Sequence (5' – 3') | Manufacturer |
|---|---|---|
| Pool_fw | CCAAGTAATACGACTCACTATAGGGCAACTCCAAGCTAGATCTACCGGT | Sigma-Aldrich |
| Pool_rev | AGTGAAAAGTTCTTCTCCTTTGCTAGCCATTTT | Sigma-Aldrich |
| Pool_oligo_rev | AGTGAAAAGTTCTTCTCCTTTGCTAGCCATTTTNNNNNNNNNNNTAGAAGCCA<br>GTAGNNNNNNNNNNNNNNNNNNNNNNNNNNNNNNACCGG<br>TAGATCTAGCTTGGAGTTGCCC | Microsynth |
| Yeast-HR_fw | CTCGTCATTGTTCTCGTTCCCTTTCTTCCTTGTTTCTTTTTCTGCACAATATTT<br>CAAGCTATACCAAGCATACAATCAACTCCAAGCTAGATCTACCGGT | Sigma-Aldrich |
| Yeast-HR_rev | CACCCTCTCCACTGACAGAAAATTTGTGCCCATTAACATCACCATCTAATTC<br>AACAAGAATTGGGACAACTCCAGTGAAAAGTTCTTCTCCTTTGCTAGC | Sigma-Aldrich |
| NGS-index_S1 | CCTT**CGTAGCTT**GGGCAACTCCAAGCTAGATCTACCGGT | Sigma-Aldrich |
| NGS-index_S2 | CCTT**CGTGCCAA**GGGCAACTCCAAGCTAGATCTACCGGT | Sigma-Aldrich |
| NGS-index_S3 | CCTT**CTAGTGCG**GGGCAACTCCAAGCTAGATCTACCGGT | Sigma-Aldrich |
| NGS-index_S4 | CCTT**CTCAGCCT**GGGCAACTCCAAGCTAGATCTACCGGT | Sigma-Aldrich |
| NGS-index_S5 | CCTT**CTCTGACA**GGGCAACTCCAAGCTAGATCTACCGGT | Sigma-Aldrich |
| NGS-index_S6 | CCTT**CTCTTCTA**GGGCAACTCCAAGCTAGATCTACCGGT | Sigma-Aldrich |
| NGS-index_S7 | CCTT**CTGATGAA**GGGCAACTCCAAGCTAGATCTACCGGT | Sigma-Aldrich |
| NGS-index_S8 | CCTT**CTGTATTC**GGGCAACTCCAAGCTAGATCTACCGGT | Sigma-Aldrich |
| NGS-index_S9 | CCTT**CTTCGCTC**GGGCAACTCCAAGCTAGATCTACCGGT | Sigma-Aldrich |
| NGS-index_S10 | CCTT**GAACTCAA**GGGCAACTCCAAGCTAGATCTACCGGT | Sigma-Aldrich |
| NGS-index_C1 | CCTT**CGAGAGCA**GGGCAACTCCAAGCTAGATCTACCGGT | Sigma-Aldrich |
| NGS-index_C2 | CCTT**CGCCAGTA**GGGCAACTCCAAGCTAGATCTACCGGT | Sigma-Aldrich |
| NGS-index_C3 | CCTT**CGCCTCAA**GGGCAACTCCAAGCTAGATCTACCGGT | Sigma-Aldrich |
| NGS-index_C4 | CCTT**CGCGGAAT**GGGCAACTCCAAGCTAGATCTACCGGT | Sigma-Aldrich |
| NGS-index_C5 | CCTT**CGCGTTAC**GGGCAACTCCAAGCTAGATCTACCGGT | Sigma-Aldrich |
| NGS-index_C6 | CCTT**CGGAAGGA**GGGCAACTCCAAGCTAGATCTACCGGT | Sigma-Aldrich |
| NGS-index_C7 | CCTT**CGGTGAGA**GGGCAACTCCAAGCTAGATCTACCGGT | Sigma-Aldrich |
| G12_fw | CCGGTAATTTAGGTCATTGAGTGAACAGACGTCAAGCTAGTTGGGCTACTG<br>GCTTCTAAAACACAGGTAAAATGG | Sigma-Aldrich |
| G12_rev | CTAGCCATTTTACCTGTGTTTTAGAAGCCAGTAGCCCAACTAGCTTGACGTC<br>TGTTCACTCAATGACCTAAATTA | Sigma-Aldrich |
| G12_M1_fw | CCGGTAATTTAGGTCATTGAGTGAACAGACGTCAAGCTAGTTGGGCTACTG<br>GCTTCTAAATTACAGGTAAAATGG | Sigma-Aldrich |
| G12_M1_rev | CTAGCCATTTTACCTGTAATTTAGAAGCCAGTAGCCCAACTAGCTTGACGTC<br>TGTTCACTCAATGACCTAAATTA | Sigma-Aldrich |
| G12_M2_fw | CCGGTAATTTAGGTCATTGAGTGAACAGACGTCAAGCTAGTTGGGCTACTG<br>GCTTCTAAAACACCGGTAAAATGG | Sigma-Aldrich |
| G12_M2_rev | CTAGCCATTTTACCGGTGTTTTAGAAGCCAGTAGCCCAACTAGCTTGACGTC<br>TGTTCACTCAATGACCTAAATTA | Sigma-Aldrich |
| G12_M3_fw | CCGGTCAATAGGTCATTGAGTGAACAGACGTCAAGCTAGTTGGGCTACTGG<br>CTTCTATTGAAAAATGG | Sigma-Aldrich |
| G12_M3_rev | CTAGCCATTTTTCAATAGAAGCCAGTAGCCCAACTAGCTTGACGTCTGTTCA<br>CTCAATGACCTATTGA | Sigma-Aldrich |
| G12_M4_fw | CCGGTAATTTAGGTCATTGAGTGAACAGACGTCAAGCTAGTAAAGACTGGC<br>TTCTAAAACACAGGTAAAATGG | Sigma-Aldrich |
| G12_M4_rev | CTAGCCATTTTACCTGTGTTTTAGAAGCCAGTCTTTACTAGCTTGACGTCTG<br>TTCACTCAATGACCTAAATTA | Sigma-Aldrich |
| G12_M5_fw | CCGGTAATTTAGGTCATTGAGTGAACAGACGTCAAGCTAGTTTCGACTGGC<br>TTCTAAAACACAGGTAAAATGG | Sigma-Aldrich |
| G12_M5_rev | CTAGCCATTTTACCTGTGTTTTAGAAGCCAGTCGAAACTAGCTTGACGTCTG<br>TTCACTCAATGACCTAAATTA | Sigma-Aldrich |
| G12_M6_fw | CCGGTAATTTAGGTCATTGAGTGAACAGACGTCAAGCAGTTGGGCTACTGC<br>TTCTAAAACACAGGTAAAATGG | Sigma-Aldrich |
| G12_M6_rev | CTAGCCATTTTACCTGTGTTTTAGAAGCAGTAGCCCAACTGCTTGACGTCTG<br>TTCACTCAATGACCTAAATTA | Sigma-Aldrich |
| G12_M7_fw | CCGGTAATTTAGGTCATTGAGTGAACAGACGTCAGCAGTTGGGCTACTGCT<br>CTAAAACACAGGTAAAATGG | Sigma-Aldrich |

| | | |
|---|---|---|
| G12_M7_rev | CTAGCCATTTTACCTGTGTTTTAGAGCAGTAGCCCAACTGCTGACGTCTGTT CACTCAATGACCTAAATTA | Sigma-Aldrich |
| G12_M8_fw | CCGGTAATTTAGGTCATTGAGTGAACAGACGTCAGCGTTGGGCTACGCTCT AAAACACAGGTAAAATGG | Sigma-Aldrich |
| G12_M8_rev | CTAGCCATTTTACCTGTGTTTTAGAGCGTAGCCCAACGCTGACGTCTGTTCA CTCAATGACCTAAATTA | Sigma-Aldrich |
| G12_M9_fw | CCGGTAATTTAGGTCATTGAGTGAACAGACGTCAACGTAGTTGGGCTACTG GCTTCTAAAACACAGGTAAAATGG | Sigma-Aldrich |
| G12_M9_rev | CTAGCCATTTTACCTGTGTTTTAGAAGCCAGTAGCCCAACTACGTTGACGTC TGTTCACTCAATGACCTAAATTA | Sigma-Aldrich |
| G12_M10_fw | CCGGTAATTTAGGTCATTGAGTGAACAGACGTCAACGTAGTTGGGCTACTG CGTTCTAAAACACAGGTAAAATGG | Sigma-Aldrich |
| G12_M10_rev | CTAGCCATTTTACCTGTGTTTTAGAACGCAGTAGCCCAACTACGTTGACGTC TGTTCACTCAATGACCTAAATTA | Sigma-Aldrich |
| G12_M11_fw | CCGGTCAATAGGTCATTGAGTGAACAGACGTCAGACCGATTGGGCTATCGG TCTCTATTGAAAAATGG | Sigma-Aldrich |
| G12_M11_rev | CTAGCCATTTTTCAATAGAGACCGATAGCCCAATCGGTCTGACGTCTGTTCA CTCAATGACCTATTGA | Sigma-Aldrich |
| G12_M12_fw | CCGGTCAATAGGTCATTGAGTGAACAGACGTCAAGCTACGTTGGGCTACGT GGCTTCTATTGAAAAATGG | Sigma-Aldrich |
| G12_M12_rev | CTAGCCATTTTTCAATAGAAGCCACGTAGCCCAACGTAGCTTGACGTCTGTT CACTCAATGACCTATTGA | Sigma-Aldrich |
| G12_M13_fw | CCGGTCAATAGGTCATTGAGTGAACAGACGTCAAGCTACAGTTGGGCTACT GTGGCTTCTATTGAAAAATGG | Sigma-Aldrich |
| G12_M13_rev | CTAGCCATTTTTCAATAGAAGCCACAGTAGCCCAACTGTAGCTTGACGTCTG TTCACTCAATGACCTATTGA | Sigma-Aldrich |
| G12_M14_fw | CCGGTCAATAGGTCATTGAGTGAACAGACAAAAAGCTAGTTGGGCTACTGG CTTCTATTGAAAAATGG | Sigma-Aldrich |
| G12_M14_rev | CTAGCCATTTTTCAATAGAAGCCAGTAGCCCAACTAGCTTTTTGTCTGTTCA CTCAATGACCTATTGA | Sigma-Aldrich |
| G12_M15_fw | CCGGTCAATAGCTGATTGAGTGAACACAGGTCAAGCTAGTTGGGCTACTGG CTTCTATTGAAAAATGG | Sigma-Aldrich |
| G12_M15_rev | CTAGCCATTTTTCAATAGAAGCCAGTAGCCCAACTAGCTTGACCTGTGTTCA CTCAATCAGCTATTGA | Sigma-Aldrich |
| G12_M16_fw | CCGGTCAATAGGACATTGAGTGAACAGACGTCAAGCTAGTTGGGCTACTGG CTTCTATTGAAAAATGG | Sigma-Aldrich |
| G12_M16_rev | CTAGCCATTTTTCAATAGAAGCCAGTAGCCCAACTAGCTTGACGTCTGTTCA CTCAATGTCCTATTGA | Sigma-Aldrich |
| G12_M17_fw | CCGGTCAATAGGACATTGAGTGAACAGTCGTCAAGCTAGTTGGGCTACTGG CTTCTATTGAAAAATGG | Sigma-Aldrich |
| G12_M17_rev | CTAGCCATTTTTCAATAGAAGCCAGTAGCCCAACTAGCTTGACGACTGTTCA CTCAATGTCCTATTGA | Sigma-Aldrich |
| G12_M18_fw | CCGGTCAATAGGAGATTGAGTGAACACACGTCAAGCTAGTTGGGCTACTGG CTTCTATTGAAAAATGG | Sigma-Aldrich |
| G12_M18_rev | CTAGCCATTTTTCAATAGAAGCCAGTAGCCCAACTAGCTTGACGTGTGTTCA CTCAATCTCCTATTGA | Sigma-Aldrich |
| G12_M19_fw | CCGGTCAATAGGAGATTGAGTGAACACTCGTCAAGCTAGTTGGGCTACTGG CTTCTATTGAAAAATGG | Sigma-Aldrich |
| G12_M19_rev | CTAGCCATTTTTCAATAGAAGCCAGTAGCCCAACTAGCTTGACGAGTGTTCA CTCAATCTCCTATTGA | Sigma-Aldrich |
| G12_M20_fw | CCGGTCAATAGGTCCAAGAGTGAACAGACGTCAAGCTAGTTGGGCTACTGG CTTCTATTGAAAAATGG | Sigma-Aldrich |
| G12_M20_rev | CTAGCCATTTTTCAATAGAAGCCAGTAGCCCAACTAGCTTGACGTCTGTTCA CTCTTGGACCTATTGA | Sigma-Aldrich |
| G12_M21_fw | CCGGTCAATAGGTCATTCAATGAACAGACGTCAAGCTAGTTGGGCTACTGG CTTCTATTGAAAAATGG | Sigma-Aldrich |
| G12_M21_rev | CTAGCCATTTTTCAATAGAAGCCAGTAGCCCAACTAGCTTGACGTCTGTTCA TTGAATGACCTATTGA | Sigma-Aldrich |
| G12_M22_fw | CCGGTCAATAGGTCATTGAGCAAACAGACGTCAAGCTAGTTGGGCTACTGG CTTCTATTGAAAAATGG | Sigma-Aldrich |
| G12_M22_rev | CTAGCCATTTTTCAATAGAAGCCAGTAGCCCAACTAGCTTGACGTCTGTTTG CTCAATGACCTATTGA | Sigma-Aldrich |
| G12_M23_fw | CCGGTCAATAGGTCATTGAGTGACAAGACGTCAAGCTAGTTGGGCTACTGG CTTCTATTGAAAAATGG | Sigma-Aldrich |
| G12_M23_rev | CTAGCCATTTTTCAATAGAAGCCAGTAGCCCAACTAGCTTGACGTCTTGTCA CTCAATGACCTATTGA | Sigma-Aldrich |
| G12_S1_fw | CCGGTCAACGGGTCATTGAGTGAACAGACGTCAAGCTAGTTGGGCTACTG GCTTCCGTTGAAAAATGG | Sigma-Aldrich |
| G12_S1_rev | CTAGCCATTTTTCAACGGAAGCCAGTAGCCCAACTAGCTTGACGTCTGTTCA CTCAATGACCCGTTGA | Sigma-Aldrich |
| G12_S2_fw | CCGGTCGATAGGTCATTGAGTGAACAGACGTCAAGCTAGTTGGGCTACTGG CTTCTATCGAAAAATGG | Sigma-Aldrich |

| | | |
|---|---|---|
| G12_S2_rev | CTAGCCATTTTTCGATAGAAGCCAGTAGCCCAACTAGCTTGACGTCTGTTCACTCAATGACCTATCGA | Sigma-Aldrich |
| G12_S3_fw | CCGGTCGCTAGGTCATTGAGTGAACAGACGTCAAGCTAGTTGGGCTACTGGCTTCTAGCGAAAAATGG | Sigma-Aldrich |
| G12_S3_rev | CTAGCCATTTTTCGCTAGAAGCCAGTAGCCCAACTAGCTTGACGTCTGTTCACTCAATGACCTAGCGA | Sigma-Aldrich |
| G12_S4_fw | CCGGTCGCTAGGTCATTGAGTGAACAGACGTCAAGCTAGTTGGGCTACTGGCTTCTAGTGAAAAATGG | Sigma-Aldrich |
| G12_S4_rev | CTAGCCATTTTTCACTAGAAGCCAGTAGCCCAACTAGCTTGACGTCTGTTCACTCAATGACCTAGCGA | Sigma-Aldrich |
| G12_S5_fw | CCGGTCGATAGGTCATTGAGTGAACAGACGTCAAGCTAGTTGGGCTACTGGCTTCTATTGAAAAATGG | Sigma-Aldrich |
| G12_S5_rev | CTAGCCATTTTTCAATAGAAGCCAGTAGCCCAACTAGCTTGACGTCTGTTCACTCAATGACCTATCGA | Sigma-Aldrich |
| G12_M3_random-template | CAGTGAAAAGTTCTTCTCCTTTGCTAGCCATTTTTCAATAGAAGCCAGTAGCCCAACTAGCTTGACGTCTGTTCACTCAATGACCTATTGACCGGTAGATCTAGCTTGGAGTT | Microsynth |
| Tetracyc_fw | CCGGTCGGCCTAAAACATACCAGATCGCCACCCGCGCTTTAATCTGGAGAGGTGAAGAATACGACCACCTAGGCCAAAATGG | Sigma-Aldrich |
| Tetracyc_rev | CTAGCCATTTTGGCCTAGGTGGTCGTATTCTTCACCTCTCCAGATTAAAGCGCGGGTGGCGATCTGGTATGTTTTAGGCCGA | Sigma-Aldrich |
| Tc-G6_fw | CCGGTCAGTAATCTGCAAACATACCAGATCGCCACCCGCGCTTTAATCTGGAGAGGTGAAGAATACGACCACCGCAGATTACTAAAATGG | Sigma-Aldrich |
| Tc-G6_rev | CTAGCCATTTTAGTAATCTGCGGTGGTCGTATTCTTCACCTCTCCAGATTAAAGCGCGGGTGGCGATCTGGTATGTTTGCAGATTACTGA | Sigma-Aldrich |
| Neo_M4_fw | CCGGTCCGGCATAGCTTGTCCTTTAATGGTCCTATGTCGAAAATGG | Sigma-Aldrich |
| Neo_M4_rev | CTAGCCATTTTCGACATAGGACCATTAAAGGACAAGCTATGCCGGA | Sigma-Aldrich |
| Tobra_fw | CCGGTGTTTCGGAAAGTTGGACCTACTGGCTTCTACCGAGCACTTAAAATGG | Sigma-Aldrich |
| Tobra_rev | CTAGCCATTTTAAGTGCTCGGTAGAAGCCAGTAGGTCCAACTTTCCGAAACA | Sigma-Aldrich |
| Tobra_N6-G5_fw | CCGGTGTTTCGGAAACTGCGGTTCTACCGAGCACTAAAATGG | Sigma-Aldrich |
| Tobra_N6-G5_rev | CTAGCCATTTTAGTGCTCGGTAGAACCGCAGTTTCCGAAACA | Sigma-Aldrich |
| Dox1_splice_fw | CGGAAGGTCTCCAGTGTTTCTCTTTTTTTCAGGAACTGGTCATTGAGTGAACAGACGTCAAGCTAGTTGG | Sigma-Aldrich |
| Dox1_splice_rev | GCCTTGGTCTCTTGGTTGTTTGGTTGTTTCAGGAACTGAAGCCAGTAGCCCAACTAGCTTGACGTCTG | Sigma-Aldrich |
